# Functional characterization of specialized immune cells in a cnidarian reveals an ancestral antiviral program

**DOI:** 10.1101/2025.01.24.634691

**Authors:** Itamar Kozlovski, Ton Sharoni, Shani Levy, Adrian Jaimes-Becerra, Shani Talice, Hee-Jin Kwak, Daria Aleshkina, Reuven Aharoni, Xavier Grau-Bové, Ola Karmi, Benyamin Rosental, Arnau Sebe-Pedros, Yehu Moran

## Abstract

Examining early-branching animal phyla can help reconstructing the evolutionary origins of immune cells. Here, we characterized immune-related cell programs in embryos of the sea anemone *Nematostella vectensis*, a model of Cnidaria, which diverged ∼600 million years ago from other animals. Using a transgenic *Nematostella* reporter line expressing mCherry under the RLRb antiviral promoter, we identified a morphologically and transcriptomically distinct cell population activated by the viral mimic poly(I:C). These cells upregulate immune effector and regulator genes and show increased phagocytic activity. Bulk RNA sequencing of RLRb expressing cells and single-cell transcriptomics revealed gene regulatory programs expressed in specialized immune cells under basal conditions and upon activation. Comparing the *Nematostella* immune expression profile with that of stony corals treated with the immunostimulant 2ʹ3ʹ-Cyclic GMP-AMP demonstrated a conserved immune response across Hexacorallia. This study uncovers a novel cnidarian immune cell type involved in antiviral immunity, providing insights into the evolutionary history of innate immunity.

## Introduction

In a world full of pathogens, organisms have been forced to evolve strategies to survive. Viruses, which infect hosts across all domains of life, play a pivotal role in driving evolution by shaping genetic diversity, facilitating horizontal gene transfer, and exerting selective pressures that influence the co-evolution of hosts and their immune systems (*1–4*). As obligate parasites, viruses rely entirely on hosts for their replication and spread, leading them to evolve strategies to evade immune detection, such as altering surface proteins, mimicking host molecules, or suppressing immune responses (*5–7*). In turn, hosts continuously adapt through innovations in innate and adaptive immunity (*8, 9*), driven by the high mutation rates and rapid evolution of pathogens (*10*). This ongoing evolutionary arms race has profoundly shaped the biological complexity and diversity of life (*11*).

Innate immune pathways are evolutionarily conserved and can be traced back to some of the most basal forms of multicellular eukaryotes. For instance, first described in the fruit fly *Drosophila melanogaster* (*12*), toll-like receptors (TLRs) and Nuclear Factor κB (NF-κB) transcription factors play a central role in anti-microbial responses across both invertebrates, as ancient as sponges (*13*) and cnidarians (*14*), and vertebrates (*15, 16*). However, until now, the recognition mechanisms and modes of action of antiviral systems have primarily been studied in vertebrates, insects, and nematodes (*17*). This limited phylogenetic coverage makes it impossible to determine the original mechanisms of these systems and the cell types involved in antiviral immunity in the last common ancestor of Bilateria. To gain new insights into the early evolution of this essential system and its components, we comprehensively investigated it in an outgroup: the sea anemone *Nematostella vectensis* (*18*), a representative model species of Cnidaria, a phylum that diverged approximately 600 million years ago from other animals (*19*). In addition to its crucial phylogenetic position, *Nematostella* is a manageable lab model with advanced molecular and gene manipulation tools, making it an excellent comparative model.

In vertebrates, antiviral responses are initiated by the recognition of double-stranded RNA (dsRNA) through pattern recognition receptors (PRRs) such as RIG-I and MDA5 (*20–22*). Upon activation, these RIG-I-like receptors (RLRs) trigger a signaling cascade via MAVS, leading to the induction of the major cytokine family of type I interferons (IFN-I) and downstream interferon-stimulated genes (ISGs) encoding proteins with antiviral functions, including Viperin, OAS, and RNase L (*23, 24*). We have previously showed that *Nematostella*, which lacks interferons, mounts an IFN-like response to dsRNA via the RLRb receptor, suggesting that this antiviral mechanism predates the evolution of interferons (*17, 25, 26*).

While much is known about vertebrate immune signaling, the origins and evolution of immune cell types remain poorly understood due to limited phylogenetic sampling, difficulties in their isolation, lack of cell type specific markers, and reliance on phenotypic similarities while neglecting gene expression profiles. Cell types are the basic functional units of multicellular organisms, still the definition of a cell type is a matter of a debate, with some definitions focusing on structure and function, while others define cell types based on specific gene expression patterns (*27, 28*). In recent years, single cell RNA sequencing (scRNAseq) technologies have generated gene expression profiles for individual cells within complex samples, enabling the identification of cell type-specific marker genes (*29, 30*). This led to the discovery of cell type subsets that were previously overlooked. For instance, scRNAseq of human blood revealed novel subsets of dendritic cells, monocytes, and progenitors (*31*).

Some invertebrates, including insects, mollusks, echinoderms, and crustaceans, have specialized immune cells that perform phagocytosis and secrete antimicrobial peptides and/or cytotoxic proteins (*32–35*). However, it is unknown whether such cells already existed in the last common ancestor of Bilateria or represent a bilaterian evolutionary innovation (*36*). While previous studies described an increased phagocytic capacity in the sea anemones *Exaiptasia diaphana* and *N. vectensis* in the context of heat stress (*37, 38*), the transcriptomic profile of such cells and their potential response to antiviral challenge have remained unexplored. scRNAseq atlas of the stony coral *Stylophora pistillata* described two putative immune cell populations expressing ISGs, IRFs and high levels of PRRs (*39*). scRNAseq analysis of *N. vectensis* revealed two distinct transcriptomic profiles potentially associated with immune functions, which are distributed between the inner and outer cell layers (*40*). These transcriptomic studies, however, were limited to basal conditions, without any immune challenge, and did not include any functional assays. Studies of *Nematostella* and *Stylophora* at the whole organism level demonstrated a highly conserved immune gene program upon treatment with the immunostimulant 2ʹ3ʹ-Cyclic GMP-AMP, encompassing numerous proteins with predicted homology to human innate immune signaling factors and known ISGs (*41, 42*). Yet, it is unknown whether the basally detected immune expression programs are functionally linked to an immune response, whether they are restricted to a certain cell type, or represent an immune state that can be activated by different cell types. In this study, we addressed these questions by a variety of transcriptomic, functional, and microscopic techniques. We demonstrate that a distinct population of cells is committed to the antiviral immune response in *Nematostella* embryos. These cells are morphologically and transcriptomically distinct and possess a unique immune gene expression program that is conserved across the cnidarian class Hexacorallia. These results suggest that specialized immune cells already existed in ancient animals before the cnidarian-bilaterian split.

## Results

### Cells expressing high levels of RLRb are morphologically and transcriptomically distinct under basal conditions

To investigate cells expressing RLRb under basal conditions, we performed intracellular immunofluorescent staining on dissociated cells from untreated 2-day-old planulae. These cells were analyzed using high-throughput imaging cytometry (ImageStream system) (*43*) alongside conventional flow cytometry and RNA sequencing (RNAseq) of fluorescence-activated cell sorting (FACS) sorted RLRb-high (RLRb^+^) and RLRb-low (RLRb^-^) cell populations (**Fig. 1A**). As a control, we used IgG stained cells to distinguish signal from noise and gated intact cells using the nuclear marker DRAQ5, as previously described (*44*) (**fig. S1, A and B**). To assess the specificity of the RLRb antibody, we performed immunoprecipitation followed by mass spectrometry. The results showed that RLRb was enriched by approximately 4.3 orders of magnitude (over 20,000-fold) based on LFQ intensities in beads incubated with the RLRb antibody, compared to control beads incubated with total rabbit IgG antibody (**Fig. 1B**), demonstrating exceptionally high specificity. We then titrated the cells with anti-RLRb antibody to determine the optimal concentration. We used the following dilutions: 1:10000, 1:5000, and 1:1000 for anti-RLRb and IgG controls. We found that a titer of 1:1000 gave a clear signal for the RLRb and a negligible background for the IgG control (**fig. S2A**). This titer was therefore used for further studies. To further assess the specificity of the antibody we measured the response to poly(I:C) treatment in embryos at 24 hours post injection (hpi). Indeed, relative to IgG control (**fig. S2B**), we observed an increase in RLRb^+^ cells from about 17 % in NaCl control to about 40 % in poly(I:C) injected individuals (**fig. S2, C and D**).

**Fig. 1.**
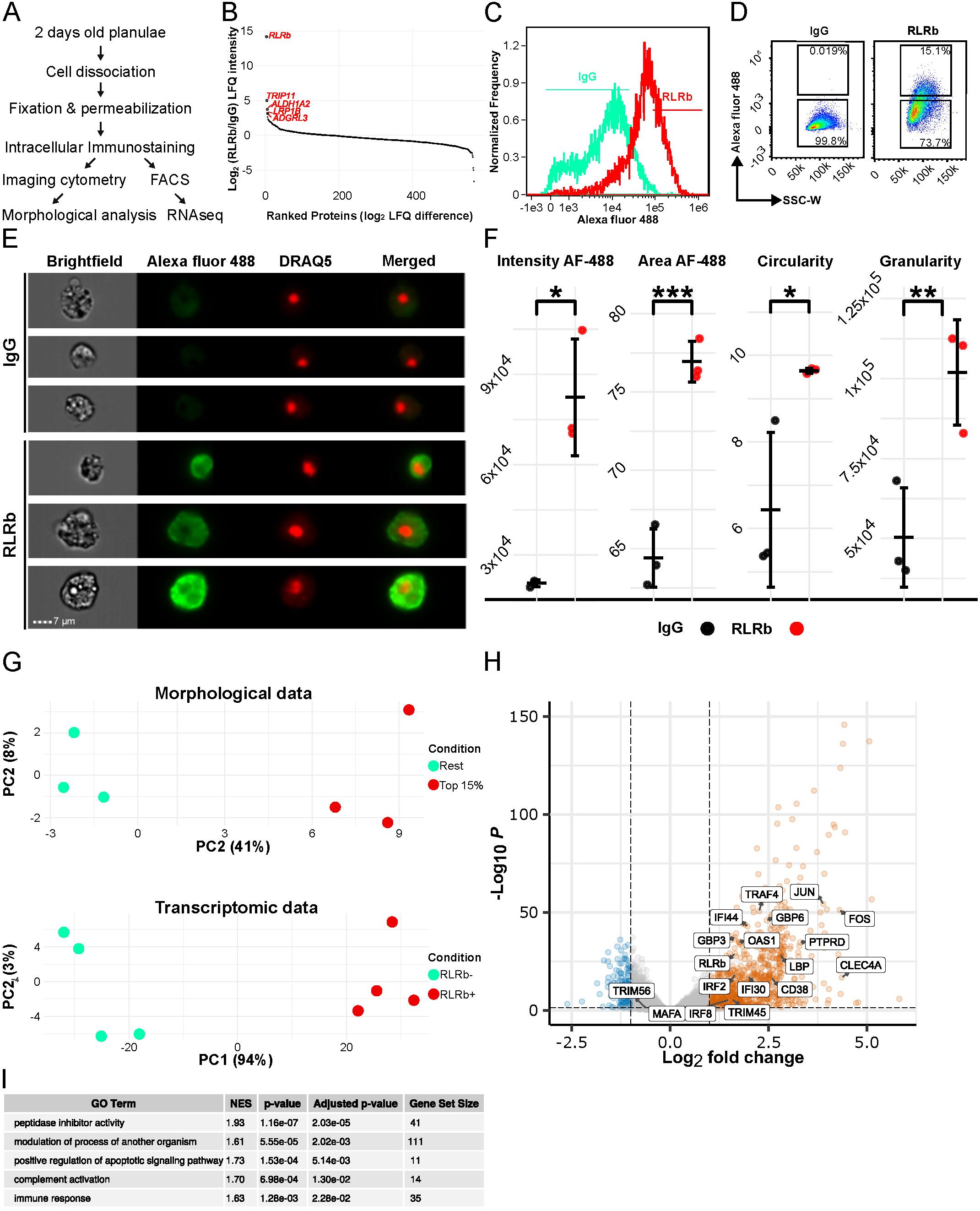
RLRb is expressed in a morphologically and transcriptomically distinct cell population under basal conditions. **(A)** Schematic overview of the experimental design. Cells were dissociated from 2-day-old planulae and subjected to intracellular immunostaining using either an RLRb-specific antibody or control rabbit IgG. **(B)** Mass spectrometry analysis of proteins immunoprecipitated with the RLRb antibody versus control IgG. RLRb is strongly enriched, confirming antibody specificity. Selected enriched proteins are labeled in red. **(C)** Representative histogram of fluorescence intensities (Alexa Fluor 488) from ImageStream analysis comparing RLRb and IgG staining. DRAQ5 was used to identify intact cells. **(D)** Representative flow cytometry analysis, as in (C). **(E)** Representative ImageStream images of cells stained with RLRb or IgG antibodies. **(F)** Morphological features of RLRb-high cells in 48-hour-old planula-derived cells. Three biological replicates were analyzed. Error bars represent standard deviation; individual data points are shown in black (IgG) or red (RLRb), with vertical lines indicating the mean. Statistical significance was assessed using a two-sided t-test: *p < 0.05, **p < 0.01, ***p < 0.001. **(G)** PCA of morphological and transcriptomic profiles comparing RLRb-high and RLRb-low cells across three and four biological replicates, respectively. **(H)** Differential gene expression analysis between RLRb-high and RLRb-low cells. Selected immune-related genes are highlighted. Differentially expressed genes are defined as those with absolute log₂ fold change > 1 and adjusted p-value < 0.05. Upregulated genes are shown in orange, downregulated in blue, and non-significant in grey. **(I)** Gene set enrichment analysis (GSEA) of differentially expressed genes. Shown are the normalized enrichment scores (NES), p-values, adjusted p-values (FDR), and the number of genes in each enriched gene set.

We then used this assay to analyze the morphological characteristics of RLRb-expressing cells under basal conditions. Cells were gated based on their RLRb signal relative to the IgG control, as determined by both imaging flow cytometry and conventional flow cytometry (**Fig. 1, C and D**). The analysis revealed that RLRb⁺ cells comprised approximately 15% of the total intact cell population, as defined relative to the IgG control (**Fig. 1D**). RLRb^+^ cells exhibited a clear cytoplasmic staining that was absent in IgG controls and a significantly higher intensity of antibody staining (**Fig. 1F)**. By global analysis of about 90,000 cells of three biological replicates, we found that RLRb^+^ cells had a significantly larger area, significantly higher circularity score, and significantly higher granularity relative to RLRb^-^ (**Fig. 1F)**. Principal component analysis (PCA) comparing 91 morphological features of the top 15 % of RLRb expressing cells against the rest of the cells in ImageStream data revealed that RLRb^+^ cells were distinct from the rest on PC1 which explains 41% of the variance (**Fig. 1G**, top panel, and **table S1**). Examination of the top loadings of PC1, the key parameters driving variability in the data, retrieved morphological features related to size such as diameter, width, and area. Next, we isolated RLRb high and low cells by fluorescence-activated cell sorting (FACS) and subjected them to RNAseq analysis. PCA analysis revealed that RLRb high cells were also transcriptomically distinct (**Fig. 1G**, bottom panel). Differential expression analysis revealed 2048 upregulated genes and 384 downregulated genes. Notably, RLRb mRNA was significantly enriched in the RLRb^+^ cell fraction (log₂ fold change = 1.65, adjusted p-value = 1.94 × 10⁻²⁹), validating the specificity of both the immunostaining and cell sorting procedures. Interestingly, among the significantly upregulated genes were genes encoding homologs of proteins involved in innate immunity such as Guanine Nucleotide-Binding Proteins (GBP3 and GBP6), 2’-5’-Oligoadenylate Synthetase 1 (OAS1), ISGs such Gamma-Interferon-Inducible Lysosomal Thiol Reductase (IFI30) and Interferon-Induced Protein (IFI44), and transcription factors such as JUN, FOS, IRF2 and IRF8 (**Fig. 1H** **and table S2**). Gene set enrichment analysis (GSEA) revealed enrichment of genes involved in complement activation (**Fig. 1G**), a process responsible for the opsonization and lytic cell death of infected cells in vertebrates (*45*). Other enriched biological processes included, positive regulation of apoptotic signaling pathway, modulation of process of another organism, peptidase inhibitor activity, and immune response (**Fig. 1I** **and table S3**). These findings highlight a population of cells with distinctive morphological and transcriptomic profiles, detectable under basal conditions, that appears to contribute to immune functions in *Nematostella*.

### RLRb is expressed in morphologically distinct cells and is upregulated in response to poly(I:C)

We previously demonstrated that RLRb mRNA and protein levels are upregulated in response to poly(I:C) exposure (*26*). However, that analysis was conducted at the whole-organism level, leaving it unclear whether RLRb expression is cell type-specific under basal conditions or following activation. To address this, we utilized a transgenic reporter line expressing mCherry under the control of the RLRb promoter, which we had previously shown to be responsive to poly(I:C) and heat stress (*46*), and designed two complementary experiments (**Fig. 2A**). First, we performed RLRb immunostaining on untreated 4-day-old wild-type planulae to assess endogenous expression under basal conditions. Second, *RLRb::mCherry* transgenic zygotes were microinjected with either poly(I:C) or NaCl as a control, followed by double immunostaining for RLRb and mCherry at 1-day post-fertilization (dpf). All samples were analyzed by confocal microscopy to visualize expression patterns at single-cell resolution.

**Fig. 2.**
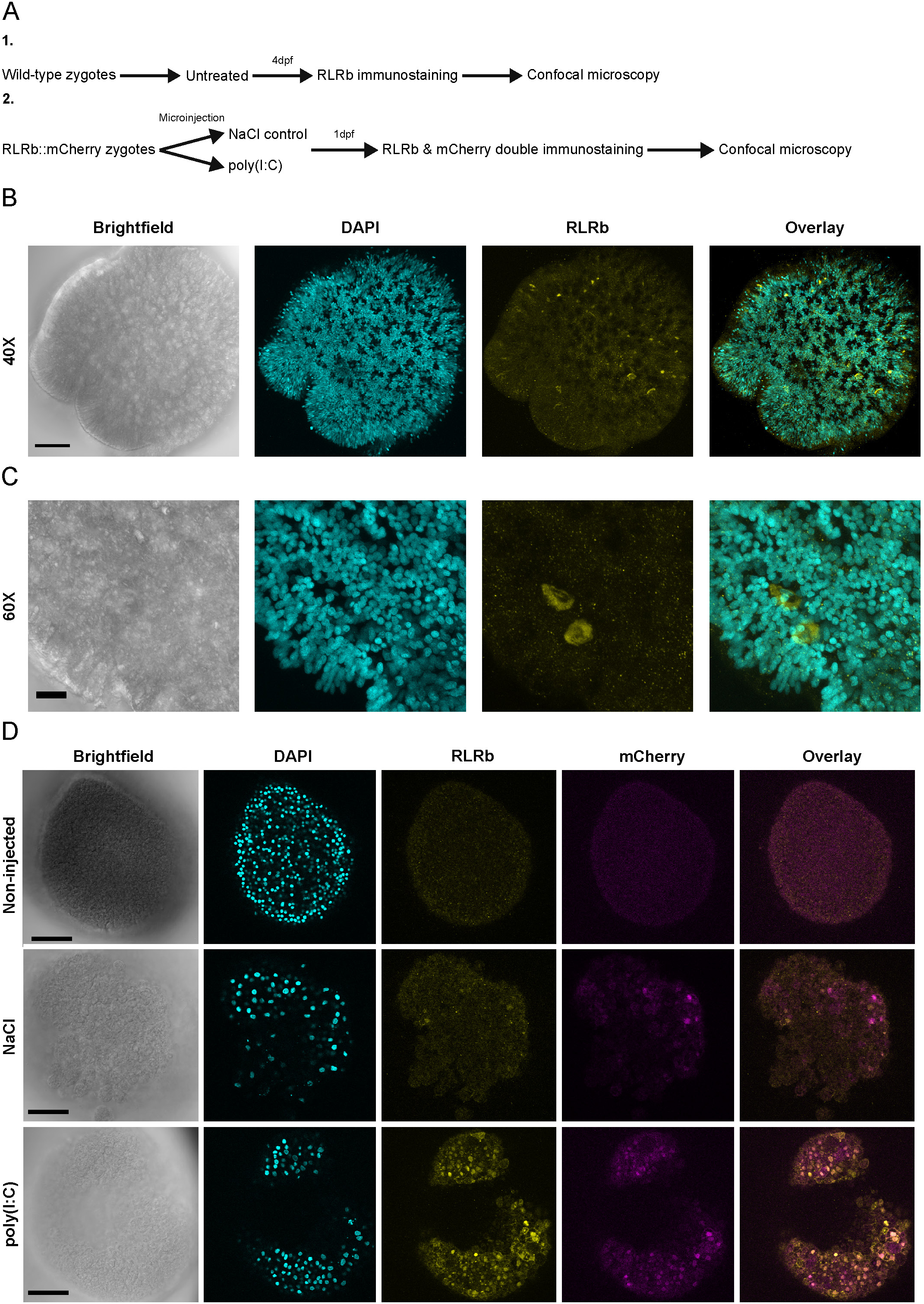
RLRb is expressed in morphologically distinct cells and is upregulated in response to poly(I:C). **(A)** Schematic overview of the experimental design. **(B)** 40× maximum intensity Z-projection confocal image of a 4-day-old planula stained with anti-RLRb, showing RLRb expression under basal conditions. Scale bar: 40 µm. **(C)** 60× maximum intensity Z-projection of a 4-day-old planula stained with anti-RLRb, highlighting cells with high RLRb expression. Scale bar: 10 µm. **(D)** Representative confocal images of non-injected control embryos. **(E)** NaCl-injected control embryos. **(F)** poly(I:C)-injected embryos. Embryos in panels (D– F) were immunostained with antibodies targeting mCherry (magenta) and RLRb (yellow). Scale bar: 50 µm.

In untreated wild-type planulae, RLRb expression was restricted to a subset of cells with a distinct morphology (Fig. 2B, C). To further examine RLRb expression and its overlap with the mCherry reporter, we performed double immunostaining on *RLRb::mCherry* embryos treated with poly(I:C), NaCl, or left untreated. Consistent with our previous western blot results (*26*), RLRb expression was low under control conditions (Fig. 2D), but strongly upregulated following poly(I:C) treatment. Likewise, mCherry expression was minimal under control conditions and increased markedly after poly(I:C) exposure. Upon stimulation, both proteins showed partially overlapping expression patterns (Fig. 2D). However, mCherry was predominantly localized in the nucleus, whereas RLRb remained mostly cytoplasmic. These findings confirm that both endogenous RLRb and the mCherry reporter are induced by poly(I:C), supporting the validity of the *RLRb::mCherry* transgenic line as a reliable reporter of RLRb activation.

### The *RLRb::mCherry* transgenic reporter line reveals a population of cells involved in antiviral immunity

Having established the specificity and responsiveness of this reporter (*46*), we next used it to explore the identity and behavior of RLRb-expressing cells during the antiviral response. To do so, we microinjected *RLRb::mCherry* transgenic zygotes with either poly(I:C) or NaCl as a control and examined mCherry expression 24 hours post-injection (hpi) using fluorescence microscopy. Embryos were then dissociated into single cells, and mCherry positive and negative populations were isolated by FACS and subjected to bulk RNAseq to characterize their transcriptional profiles (**Fig. 3A**). Similarly to our previous double immunostaining analysis (**Fig. 2D**), we observed an increase in mCherry expression upon poly(I:C) treatment relative to WT or NaCl controls detected by Fluorescent microscopy (**Fig. 3B**). This was quantified at the cellular level by flow cytometry (**Fig. 3C and fig. S3A**). On average, approximately 30 % of cells from poly(I:C) treated embryos were mCherry-positive, significantly higher compared to 8 % in NaCl-treated *RLRb::mCherry* derived cells and 1 % in their non-injected WT counterparts (**Fig. 3D)**. In agreement with our previous morphological analysis, cells within the mCherry negative fraction were relatively small (low FSC-A) and non-granular (low SSC-A) compared to mCherry positive cells (**Fig. 3E**). This difference was statistically significant with, on average, approximately 80 % of the mCherry positive cells being large and granular compared to approximately 40 % in the mCherry-negative cells (**Fig. 3F**).

**Fig. 3.**
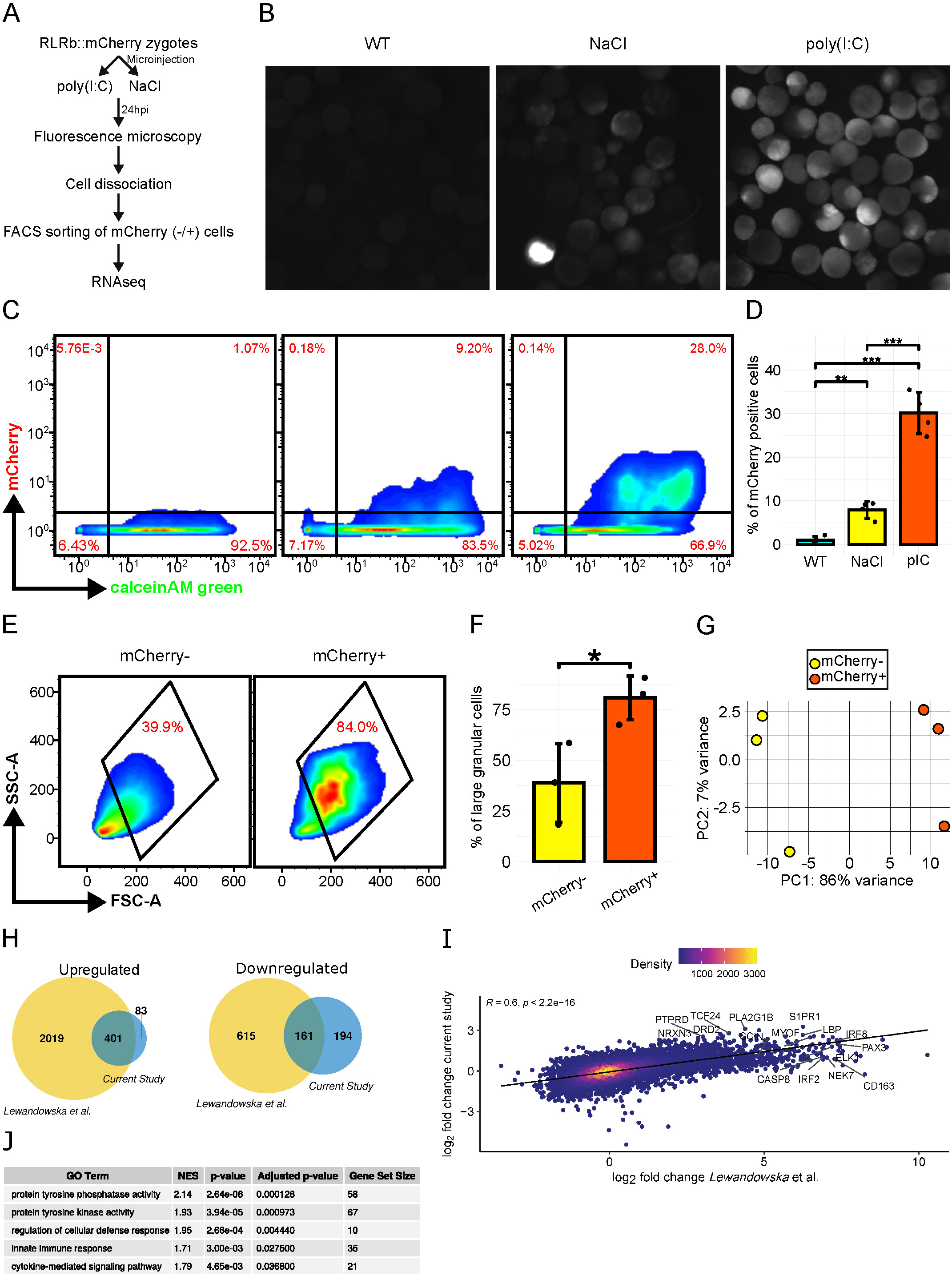
The *RLRb:mCherry* transgenic reporter line reveals morphological and transcriptomic features of dsRNA activated cells. (A) Schematic overview of the experimental design**. (B)** mCherry expression in embryos, left to right: non-injected WT control, NaCl control, and poly(I:C) injected. **(C** and **D)** Flow cytometry of cells dissociated from embryos at the same order as (A). **(E)** Forward side scatter (FSC) against side scatter (SSC) profile of mCherry negative cells **(left)** and mCherry positive cells **(right). (F)** Quantification of cells gated as shown in panel E. **(G)** PCA comparing transcriptomes of mCherry positive (mCherry+) versus negative (mCherry-) cells. **(H)** Venn diagrams comparing differentially expressed genes in the current study compared to those found by Lewandowska *et.al* (*26*). The same analysis was carried for both datasets. **(I)** Scatter plot showing the correlation of log₂ fold changes for genes identified in the current study (y-axis) and those reported by Lewandowska et al. (x-axis) (*35*). Each point represents a gene, and color intensity indicates the local point density, with warmer colors denoting higher densities. Selected genes of interest are labeled. The black regression line highlights the positive correlation between datasets (Pearson correlation coefficient R = 0.6, p < 2.2e−16 (H). **(J)** GSEA showing pathways that were enriched in mCherry positive cells. Shown are the normalized enrichment scores (NES), p-values, adjusted p-values (FDR), and the number of genes in each enriched gene set. For bar plots, mean is shown with error bars representing standard deviation. Two-sided t test was performed: * p<0.05, **p<0.01, ***p<0.001.

Next, we sorted mCherry positive and negative cells, extracted their RNA, and subjected them to RNAseq. Using 3 biological replicates. We observed a clear distinction between negative and positive cells explained by PC1, which explains 86 % of the variance in the data (**Fig. 3G**). In total, 484 upregulated genes and 355 downregulated genes were identified by differential expression analysis. Despite differences in experimental setup, comparison of these genes to a previously published study that compared the effect of poly(I:C) at the whole animal level (*26*) demonstrated a striking similarity with 401 out of 484 (∼83 %) overlap of upregulated genes and 161 out of 355 (∼45 %) overlap in downregulated genes (**Fig. 3H**). Genes with an adjusted p value of less than 0.05 from both studies were positively correlated in their log 2-fold change expression with R= 0.8 (Pearson R=0.6 for all genes) (**Fig. 3I** **and table S4**). Among the upregulated genes, GSEA revealed enrichment for cytokine mediated signaling pathway, innate immune response, protein tyrosine kinase activity, protein tyrosine phosphatase activity, and regulation of cellular defense response (**Fig. 3J** **and table S5**). Altogether, these results indicate that the antiviral immune response in *Nematostella* is activated in a certain cell population with distinctive morphological and transcriptomic profiles. Similar to vertebrates, in which mitogen-activated protein kinases (MAPKs) are activated by PRRs to promote inflammation (*47*), this cell population upregulated homologs of tyrosine kinases, receptor tyrosine kinases (RTKs), components of the MAPK signaling pathway, and phosphatases such as Protein Tyrosine Phosphatase Receptor Type D (PTPRD), FER tyrosine kinase (FER), Distinct Subgroup Of The Ras Family Member 1 (DIRAS1), Fibroblast Growth Factor Receptor 1 (FGFR1), and Rac Family Small GTPase 2 (RAC2), Mitogen-Activated Protein Kinase 8 Interacting Protein 1 (MAPK8IP1), among others.

### *RLRb::mCherry* poly(I:C) induced cells are phagocytic

The observation that a certain cell population is activated by poly(I:C) and upregulates genes belonging to the Rho family GTPases, including RAC and RAS homologs, and other genes potentially involved in phagocytosis such Fascin (FSCN1) homolog, ELMO1 homolog, and Macrophage Expressed 1 (MPEG1) (*48, 49*), together with previous studies that described phagocytic cells in Hexacorallia (*37*) prompted us to ask whether mCherry-expressing cells are capable of phagocytosis. To address this, we conducted a set of phagocytosis assays. As before, zygotes were injected with poly(I:C) or NaCl, and non-injected *RLRb::mCherry* and WT embryos were used as controls. At 24 hpf, embryos were dissociated into single cells and incubated overnight with phagocytic probes. Engulfment of the phagocytic probe was then assessed and compared between the fraction of mCherry negative and positive cells by flow cytometry (**Fig. 4A** and fig. **S4A**). As before, mCherry expression was absent in WT derived cells, nearly absent in non-injected *RLRb::mCherry* derived cells, slightly induced in NaCl controls with approximately 3 %, and induced in about 18 % of cells in the poly(I:C) treated group, in the absence of a phagocytic probe (**Fig. 4, B** and **C**). Incubation with the antigenic protein DQ ovalbumin which can only emit green fluorescence upon proteolytic degradation (*50*) was enriched in mCherry positive cells, with approximately 32 % versus about 12 % in the negative fraction (**Fig. 4, D** and **E**). Similarly, incubation with the Gram negative *Escherichia coli* pHrodo green bioparticles conjugate (*51*), which is non-fluorescent outside the cells and becomes fluorescent in phagosomes, resulted in detectable phagocytes that were mostly notable upon poly(I:C) treatment (**Fig. 4F**). The fraction of mCherry positive cells was significantly enriched for phagocytes, with about 42 % of the cells being positive versus approximately 12 %, respectively (**Fig. 4G**). Lastly, incubation with the Gram-positive *Staphylococcus aureus* pHrodo green bioparticles conjugate, which acts similarly to *E. coli* pHrodo and emits fluorescence at low pHs inside phagosomes, was enriched in mCherry positive cells, with approximately 52 % of the cells being positive versus 20 % in the mCherry negative fraction (**Fig. 4**, **H** and **I**). These results indicate that poly(I:C) activation in mCherry positive cells induces robust phagocytosis.

**Fig. 4.**
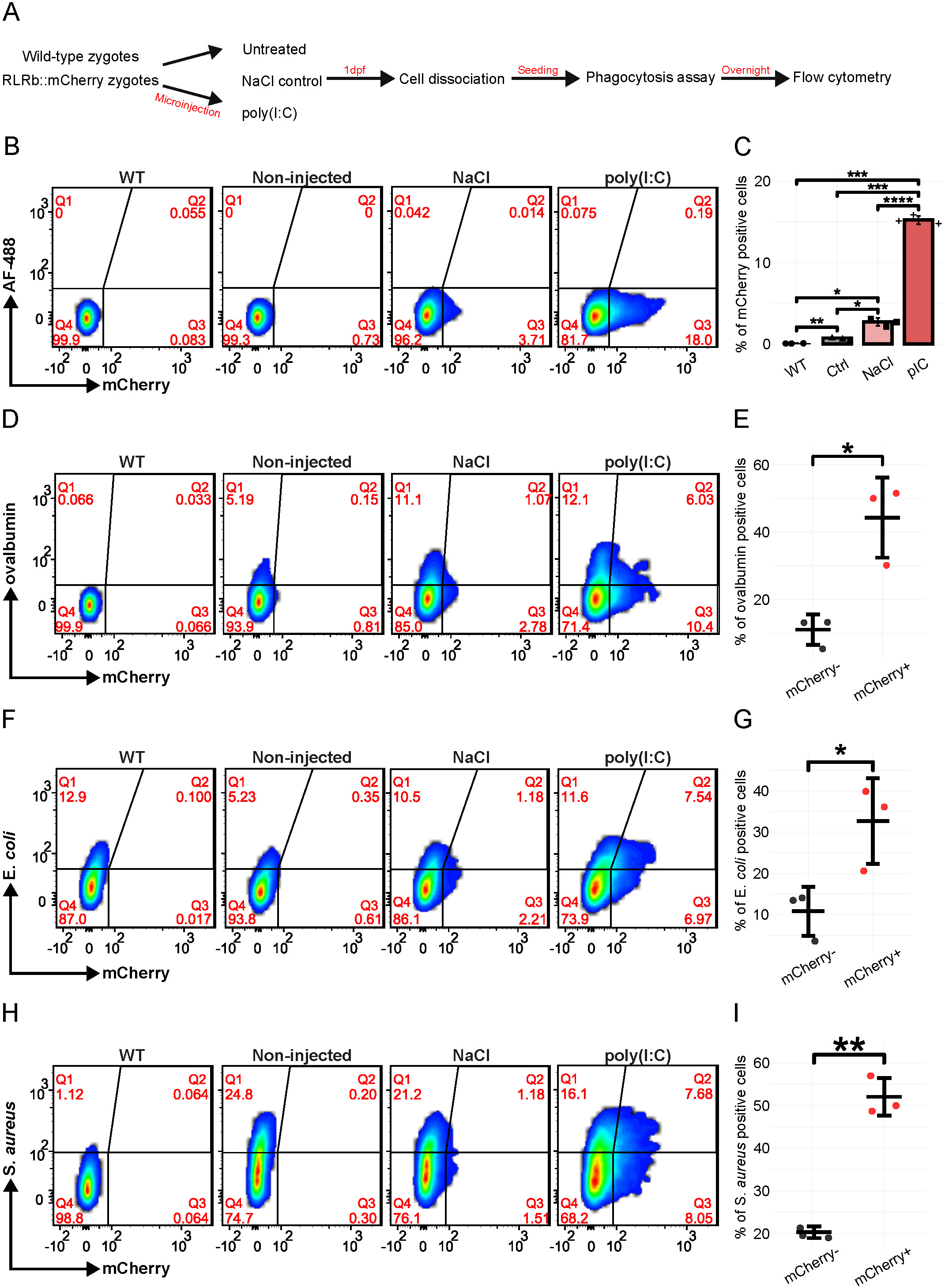
The *RLRb:mCherry* positive cells are phagocytic. Cells were dissociated at 24 hpf and incubated for 16 hours with or without phagocytic probes. **(A)** Schematic overview of the experimental design. **(B)** Representative flow cytometry results of untreated controls. **(C)** Quantification of three biological replicates as described in (B). **(D** and **E)** cells were incubated with DQ ovalbumin and the percentage of positive cells was calculated based on green probe fluorescence and mCherry expression. **(F** and **G)** Cells were incubated with E. *coli* phagocytic probe. **(H** and **I)** cells were incubated with S. *aureus* phagocytic probes. For all quantifications three biological replicates were used and average is shown. Error bars represent standard deviation and individual data points are shown. For panel (B) one way ANOVA test with Tukey’s post hoc test was performed. For panels (D,F, and H) Two-sided t test was performed: * p<0.05, **p<0.01, ***p<0.001.

### dsRNA activation is restricted to immune cells and is linked to immune gene expression modules

Elaborating on our previous observations that RLRb expression is restricted to a certain population of cells, both under basal condition and upon activation with dsRNA, we conducted scRNAseq to determine more precisely the cell type composition and gene expression programs associated with the immune response. We used two biological replicates of the *RLRb::mCherry* transgenic line, applying the same conditions as before: non-injected (Ctrl), NaCl-injected control (NaCl), and poly(I:C)-injected embryos (pIC). More than 80% of cells were viable across all treatments, with 23% and 29% of cells showing mCherry positivity in the two biological replicates of poly(I:C)-treated embryos. These mCherry fluorescence levels were consistent with those from previous experiments (**Fig. 3C**), confirming the reliability of the assay. Following dissociation, cells were fixed using the ACME method (*44*) and each condition was labeled with different ClickTag barcodes (*52*). We conducted two replicates of a multiplexed scRNAseq analysis of the three conditions together, to be able to quantitatively analyze cell type composition differences, in addition to gene expression differences in a cell type-specific manner.

In total, we obtained 11,550 single-cell transcriptional profiles that we clustered into 187 metacells (*53*). The two biological replicates, namely Nvec01 and Nvec02, exhibited no batch effects (**fig. S5**, **A** and **B**) and were, therefore, merged for further analysis. Examination of mCherry mRNA expression at single-cell level across conditions was consistent with our previous results at the protein level (**Fig. 3**, **A** to **C**), showing enrichment upon poly(I:C) treatment (**Fig. 5A**). This enrichment was positively correlated with RLRb expression levels across metacells and across cells (**Fig. 5B, fig. S5C and table S6**), R = 0.84 (Pearson correlation coefficient), supporting the fact that the *RLRb* promoter transgenic reporter line strongly recapitulates endogenous *RLRb* expression. In agreement with our previous bulk-RNAseq (**Fig. 3, J** and **K, table S7**), differential expression analysis of poly(I:C) versus NaCl treated cells revealed enrichment of immune related genes such as IRF8 and GBP7, protein tyrosine phosphatases, kinases such as PTPRD and TEK, and others (**table S7**). Strikingly, poly(I:C) derived cells formed a distinct cell cluster and a group of metacells which were nearly devoid of non-injected or NaCl derived cells (**Fig. 5D** and **fig****. S5A**). By calculating the average expression levels score of all genes that were upregulated in the mCherry positive fraction of cells (**Fig. 3I** **and table S4**), we found that this set of genes was, indeed, almost exclusively expressed in the cell cluster that was unique to poly(I:C) (**Fig. 5E**).

**Fig. 5.**
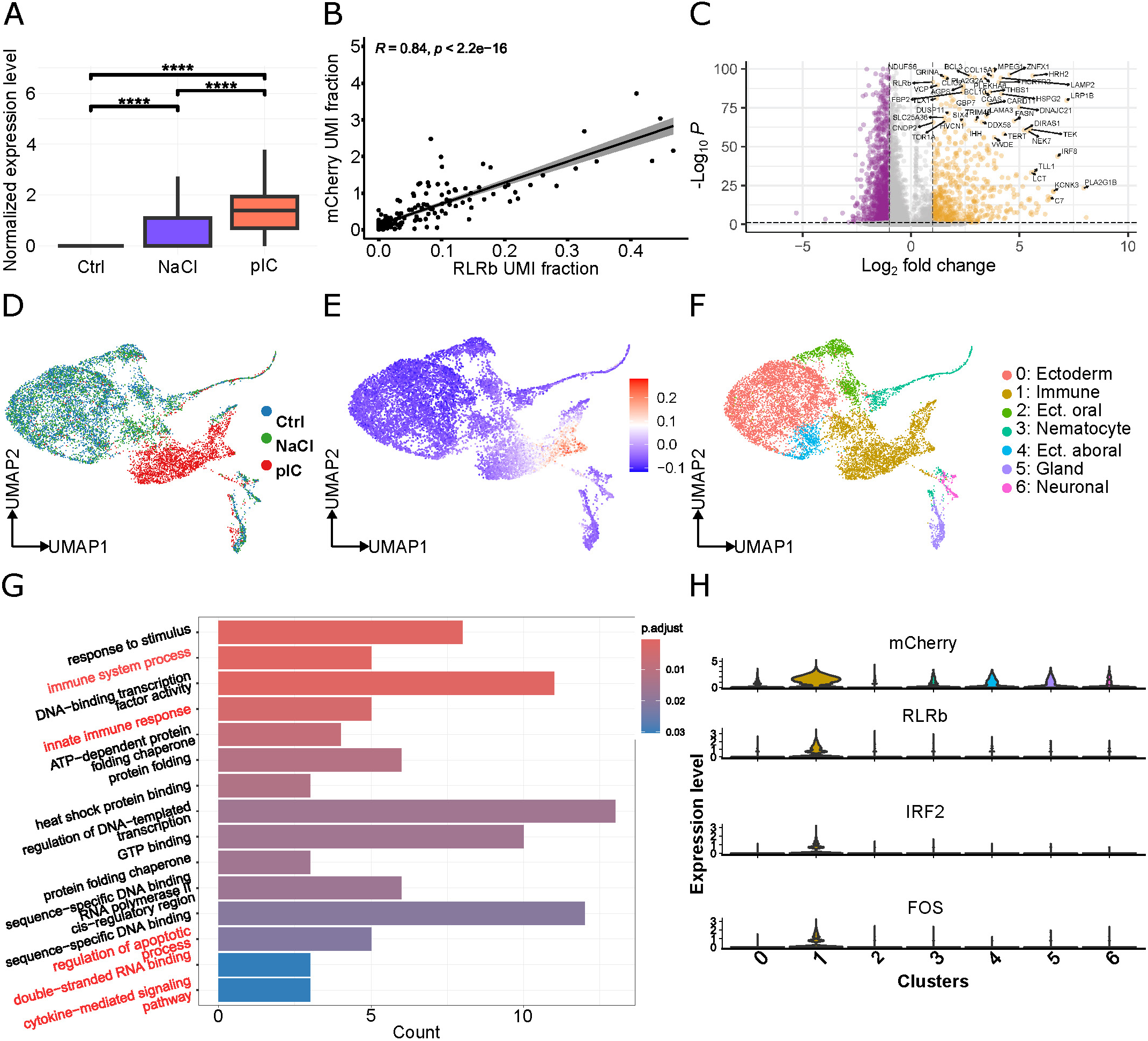
scRNAseq reveals an immune cell cluster. **(A)** normalized mRNA expression level of mCherry is shown across conditions. **(B)** Scatter plot showing the Pearson correlation of unique molecular identifier (UMI) fractions of RLRb and mCherry across metacells. **(C)** Volcano plot of differentially expressed genes using scRNAseq between poly(I:C) and NaCl cells. Genes that are significantly upregulated in poly(I:C) condition are shown in orange. Significantly downregulated genes are shown in magenta. **(D)** UMAP projection based on conditions. **(E)** Average expression score was computed for genes that were significantly upregulated in mCherry positive cells (Fig. 3I) and their expression is shown on the UMAP projection. **(F)** Clustering and annotations of the scRNAseq data. **(G)** GSEA for differentially expressed genes found in cluster 1 (immune). Top 15 categories are shown. Processes directly related to immune response are highlighted in red. **(H)** Expression levels of selected genes across different clusters.

Next we annotated cell types using previously published cell type markers (*40*) (**table S8**). Using these markers, we were able to identify gland cells, neurons, embryonic ectodermal cells, and a small subset of nematocytes (**Fig. 5F**, **fig. S6A and table S9**). The expression of IRF1-2a, which was reported as a putative immune cell marker (*40*), together with RLRa and RLRb was enriched in cluster 1 of our data (**fig. S6A**). Other genes that were strongly enriched in cluster 1 included: Radical S-Adenosyl Methionine Domain Containing 2 (RSAD2, known as viperin) a known ISG directly involved in viral inhibition (*54*), toll like receptor (TLR), GBPs as well as other ISG homologs (**table S10**). Indeed, GSEA of genes that were differentially expressed in cluster 1 revealed enrichment for immune related processes such as: innate immune response, response to stimulus, regulation of apoptosis, double-stranded RNA binding, cytokine mediated signaling pathway, and terms potentially related to the activated state such as DNA binding transcription activity, sequence specific DNA binding, and GTP binding (**Fig. 5G** **and table S10**). The mCherry gene, the gene encoding RLRb, IRF2, and FOS were expressed almost exclusively in cluster 1, further supporting its cell type identity (**Fig. 5H**).

In order to infer gene programs from our scRNA-seq data, we calculated gene modules using weighted correlation network analysis (WGCNA). We identified 19 different gene modules, termed GS1 to GS19, corresponding to different metacells (**fig. S7A and table S11**). This analysis revealed five different gene sets that were exclusive or highly enriched in cluster 1, representing associated immune programs (**fig. S7B**). ORA of individual gene modules revealed module specific significant enrichment of biological processes such as siRNA processing, double stranded RNA binding, and helicase activity (**fig. S8A and table S12**, corresponding to GS17), sterol transport and binding, adenylate cyclase-inhibiting G protein-coupled glutamate receptor signaling pathway (**fig. S8B and table S13**, corresponding to GS16), actin binding, cell differentiation, positive regulation of cell population proliferation (**fig. S8C and table S14**, corresponding to GS14), and plasma membrane organization, kinase activity, and Nf-κB binding (**fig. S8D and table S15**, corresponding to GS15). Other gene modules were unique to non-immune clusters, further supporting the specificity of our results (**fig. S7B**). By correlating our metacell results with those published by Cole *et al.* (*40*), we identified with high confidence gland cells, nematocytes, neurons, and embryonic ectodermal cells (**fig. S9A**).

### Cell composition changes dramatically upon activation and is associated with temporal changes in gene expression related to cell division

Remarkably, our analysis uncovered a significant shift in cell type composition following poly(I:C) treatment compared to controls, with the majority of cells associating with the immune cluster 1 (**Fig. 6A**). To further investigate the temporal changes in gene expression in response to dsRNA, we subclustered cluster 1 into four distinct clusters (**Fig. 6B**), each of which had distinct gene markers (**Fig. 6C** **and table S16**). Next, to investigate the dynamics of gene expression changes upon activation with poly(I:C), we conducted a pseudotime analysis using Slingshot (*55*). This revealed the starting and endpoint of the immune response activation (**Fig. 6D**). This prediction was in agreement with the cell distribution across conditions, where cluster 1 and 2 (in the subclustered data) were enriched upon poly(I:C) treatment and nearly absent in controls (**Fig. 6E**). This suggests that poly(I:C) triggers a robust shift in cell states, possibly indicating activation, proliferation, or differentiation of a subset of cells, which may represent an antiviral response. Next, we conducted over-representation analysis (ORA) for genes that were upregulated in sub-clusters 0 and 1, representing the basal state and the initial activation, respectively. Interestingly, sub-cluster 0 which was inferred to form early upon activation and was present at basal conditions, was enriched for terms related to cell cycle such as cell division, microtubule cytoskeleton organization, and RNA polymerase II cis-regulatory region sequence specific DNA binding, potentially indicating proliferation (**Fig. 6F** **and table S17**). Sub-cluster 1 which was predicted to form early in the response was enriched in terms related to cellular organization such as actin binding, protein transport, GTP binding, and endoplasmic reticulum to Golgi vesicle-mediated transport (**Fig. 6F**), potentially indicating cellular changes that might be related to phagocytosis (**Fig. 4** and table **S18**) (*56*). Sub-cluster 2 was mainly enriched for “transcription coregulator activity” (**table S19)** and sub-cluster 3 was enriched for processes related to ribosome biogenesis, GTP binding and GTPase activity **(table S20**). We modeled the expression of selected genes across the pseudotime axis (**Fig. 6G)** and found that mCherry expression, as expected, was initiated at the early activation step (cluster 1) and reached a peak in the transition to the later stage, cluster 2. Similarly, the gene encoding the antiviral GBP3, was induced early in the response, whereas the Macrophage-expressed gene 1 (MPEG1), a highly conserved gene encoding the pore forming perforin-2 protein used for antimicrobial defense in invertebrates (*57*), was induced at the late stage. Genes that were specifically expressed in cluster 3 included the mediator of apoptosis Caspase 8 Associated Protein 2 (CASP8AP2).

**Fig. 6.**
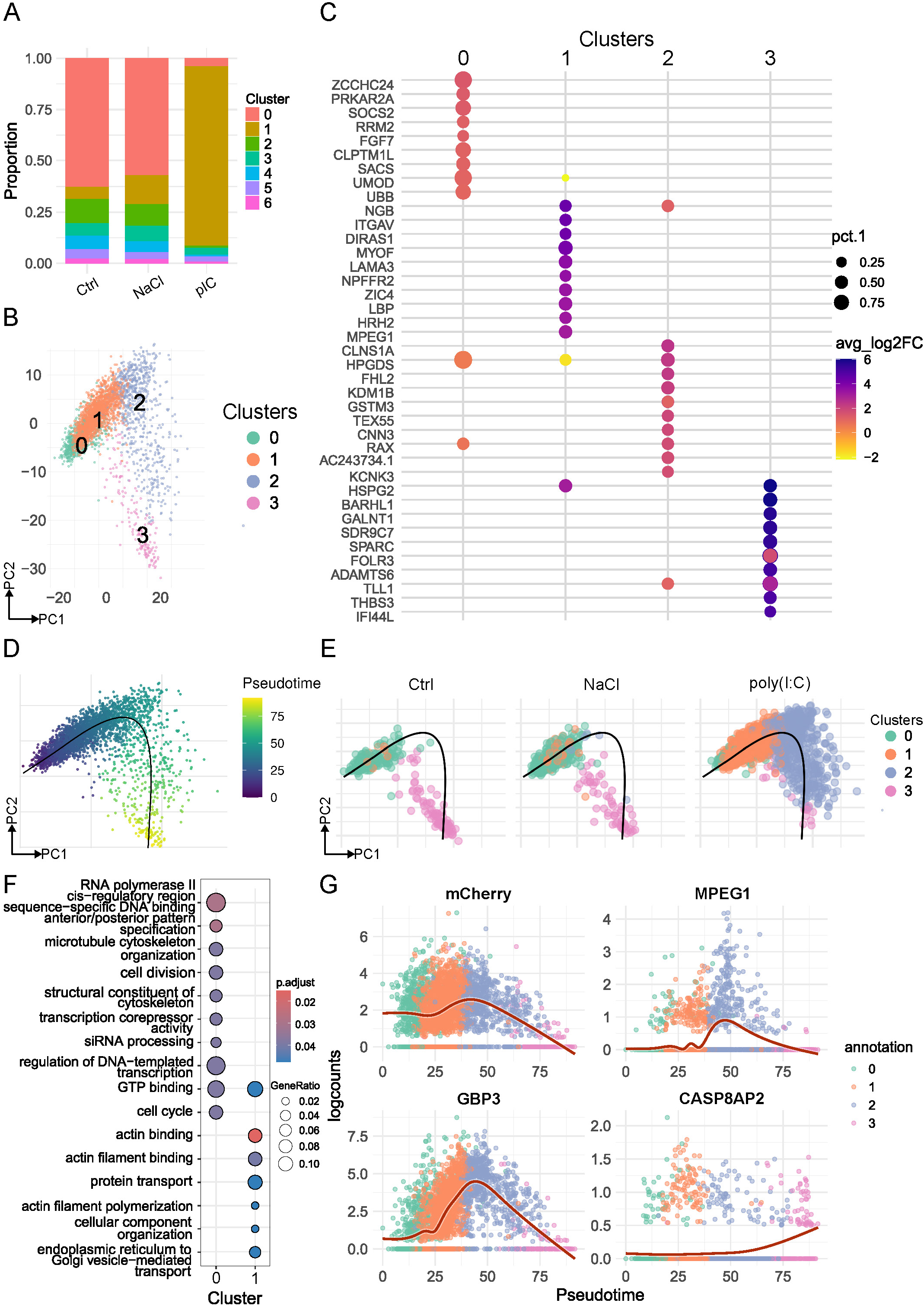
Cellular composition shifts drastically following dsRNA challenge and is accompanied by dynamic changes in gene expression over time. **(A)** Proportion of cell cluster across different conditions. **(B)** PCA suclustering of cluster 1. **(C)** Top 10 cell state markers of each cluster shown in B. **(D)** Pseudotime analysis predicts the trajectory of gene expression changes. **(E)** Pseudotime prediction as shown in D for each of the conditions. **(F)** ORA of marker genes identified in sub-cluster 0 and in sub-cluster 1 **(G). (H)** Fitted models of pseudotime-dependent gene expression of selected genes.

### Comparative analysis reveals conserved and species-specific antiviral responses

To characterize the molecular features distinguishing the activated and basal cell states within cluster 1, we performed differential expression analysis to identify key marker genes. These genes were then functionally classified into categories such as transcription factors, receptors, surface proteins, and effectors based on gene ontology annotations and homology to known genes (**Fig. 7A and fig. S10A).** In the basal (naïve) state, homologs of transcriptional regulators such as BARHL1, MAFA, DBP, and SNAI1 were enriched, along with receptors like RLRa, RLRb, FOLR3, TAAR9, and FZD7. Effector genes included known immune-related homologs such as MAVS, MIF, and OAS1. In contrast, the activated poly(I:C) state was characterized by strong upregulation of immune-associated transcription factors, including IRF2, FOS, ELF1, STAT5A, and ELK1. Predicted surface receptors such as HCRTR2, GPR157, UNC5C, and EDAR2 were also enriched. Notably, activated cells showed induction of key immune effectors including RNF213, GBP1, STING1, CASP3, and MPEG1, highlighting a transcriptional program consistent with antiviral immunity. Additional predicted transcription factors upregulated in each metacell are listed in **fig. S10**.

**Fig. 7.**
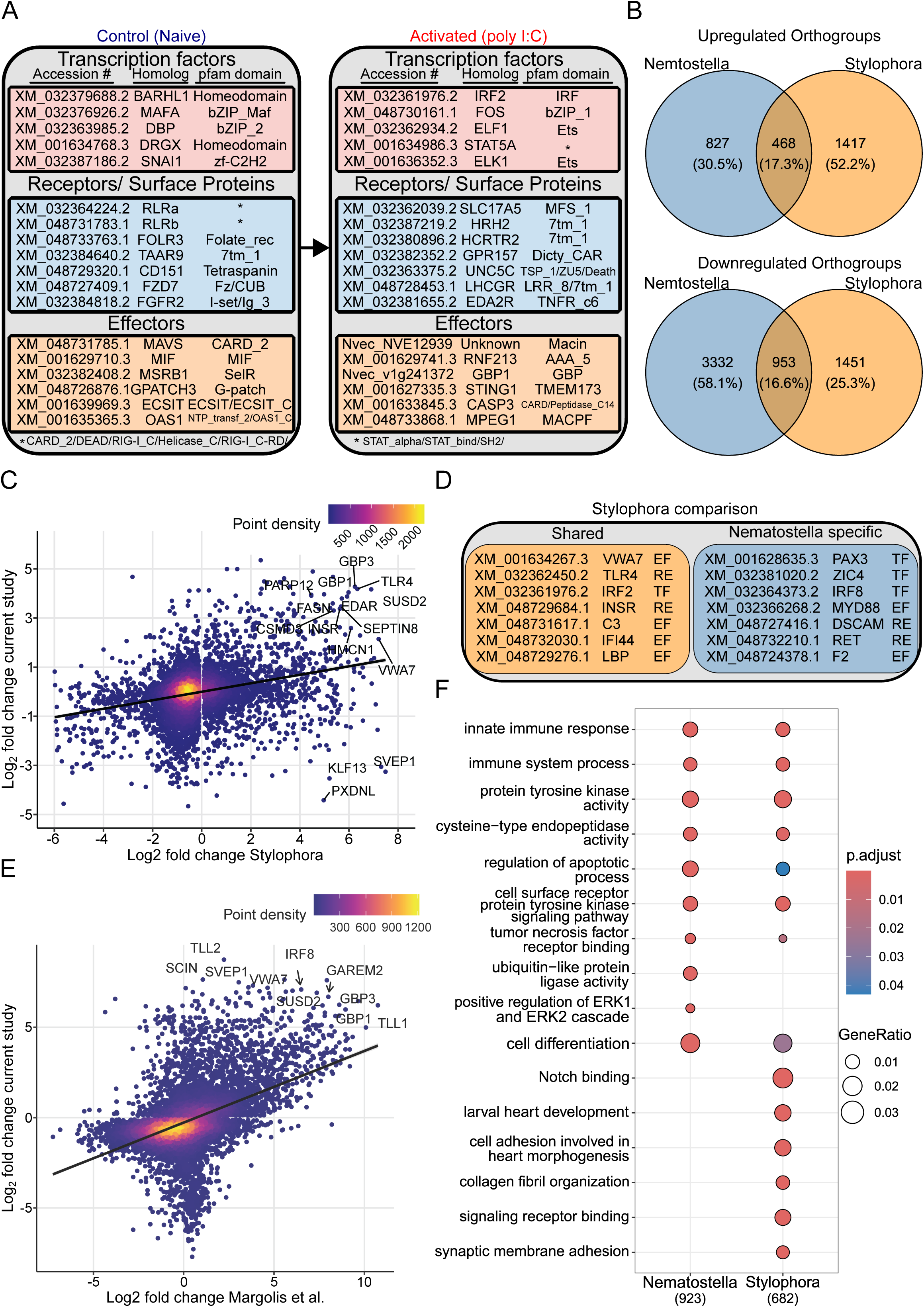
Cross-species comparison of immune gene responses between *N. vectensis* and *S. pistillata*. **(A)** Differentially expressed genes in the immune cluster of Nematostella following poly(I:C) stimulation, categorized into transcription factors (TF), receptors/surface proteins (RE), and effectors (EF). Genes specific to the naïve or activated state are shown with corresponding domain annotations. **(B)** Overlap of differentially expressed orthogroups in *Nematostella* and *Stylophora* following immune stimulation. Venn diagrams display shared and species-specific upregulated (top) and downregulated (bottom) orthogroups. cGAMP treated S. *pistillata* published by Li *et.al* (*41*) was used for comparison**. (C)** Correlation of log₂ fold changes for upregulated orthologous genes between *Stylophora* and the current *Nematostella* study. Each dot represents a gene, with color indicating local point density. Selected conserved genes are annotated**. (D)** List of shared and *Nematostella*-specific upregulated genes with known immune-related domains, grouped into transcription factors (TF), receptors (RE), and effectors (EF) **(E)** Correlation of log₂ fold changes for upregulated genes between the current study and and those identified in cGAMP treated *Nematostella* polyps published by Margolis *et.al* (*42*). **(F)** GO term enrichment of upregulated orthologous genes in *Nematostella* (left) and *Stylophora* (right). Dot size indicates gene ratio; color reflects adjusted p-value. Immune-related terms such as innate immune response and tumor necrosis factor receptor binding are enriched in both species.

To assess the generalizability of our findings to other Hexacorallia species, we compared the differentially expressed genes identified in the immune cluster to a recent study by Li *et al.* (*41*), which demonstrated that the stony coral *S. pistillata* activates a robust immune response when exposed to 2ʹ3ʹ-Cyclic GMP-AMP (cGAMP), a cyclic dinucleotide that activates the cGAS-STING pathway responsible for cytosolic DNA detection (*58*). Despite the differences in experimental design, treatment and species, comparison of orthologous genes identified using OrthoFinder (*59*) (**table S21**) revealed both shared and species-specific components of the immune response.

To evaluate this further, we compared differentially expressed orthogroups between *Nematostella* and *Stylophora* following immune stimulation. Among upregulated orthogroups, 17.3 % (468) were shared, while 30.5 % (827) and 52.2 % (1,417) were specific to *Nematostella* and *Stylophora*, respectively. For downregulated orthogroups, 16.6 % (953) were shared, with 58.1 % (3,332) unique to *Nematostella* and 25.3% (1,451) unique to *Stylophora* (Fig. 3B). These results suggest the existence of a core antiviral program shared between the species, along with substantial species-specific transcriptional signatures likely shaped by evolutionary divergence.

To explore gene-level conservation, we plotted the log₂ fold change of orthologous genes from both species (**Fig. 7C** **and table S22**). While the overall correlation was modest (Pearson R=0.40 when adjusted p value<0.05), several immune-related genes such as GBP1, GBP3, TLR4, EDAR, and IRF2 were consistently upregulated in both *Nematostella* and *Stylophora*. These genes likely represent evolutionarily conserved components of the antiviral immune response. A representative subset of these shared and *Nematostella*-specific immune genes is shown in **Fig. 7D**. Shared genes included transcription factors (IRF2), receptors (TLR4, INSR), and effectors (C3, IFI44, LBP), while *Nematostella*-specific upregulated genes included PAX3, IRF8, MYD88, and the receptors DSCAM and RET.

To assess consistency with previous findings within *Nematostella*, we compared our results to a study by Margolis *et al.* (49), in which polyps were treated with cGAMP. This comparison revealed a stronger positive correlation (Pearson R = 0.66 for adjusted p value<0.05) between the datasets (**Fig. 7E**), supporting the robustness of the antiviral transcriptional program across stimuli and developmental stages. Shared upregulated genes included canonical immune effectors such as GBP1, GBP3, IRF8, SUSD2, TLL1, and VWA7 (Tables S4 and table S23), further highlighting a conserved set of antiviral genes within *Nematostella*.

Finally, we performed Gene Ontology (GO) over-representation analysis of upregulated genes in both species to better understand the biological processes underlying the response. Both *Nematostella* and *Stylophora* showed enrichment for core immune-related terms such as "innate immune response," "immune system process," "regulation of apoptotic process," and "protein tyrosine kinase activity" (**Fig. 3F**). However, species-specific enrichments were also evident. *Stylophora* was uniquely enriched for developmental and adhesion-related terms such as "larval heart development," "collagen fibril organization," and "synaptic membrane adhesion," while *Nematostella* showed enrichment for **"**positive regulation of ERK1 and ERK2 cascade**"** and **"**ubiquitin-like protein ligase activity**."** These findings highlight both shared and divergent transcriptional programs shaping the antiviral response in Hexacorallia. Other shared processes included RNA interference (RNAi), NF-κB transcription factor binding, GTPase binding and activity. Notably, all these processes were previously indicated to play a role in cnidarian immunity (*25, 26, 60*). These similarities in immune response regulation and shared components with the identified immune gene program, point to an ancestral gene regulatory program that can be activated in immune cells of various species of Hexacorallia following different immunogenic cues.

## Discussion

Multicellular organisms rely on diverse immune defense strategies, often employing multiple mechanisms simultaneously to ensure survival. Among these, vertebrates have evolved a highly advanced adaptive immune system, characterized by an almost limitless variety of antigen receptors that are clonally expressed on specialized immune cells called lymphocytes (B and T cells) circulating throughout the body and detecting pathogens (*61*). Long before the adaptive immune system evolved, organisms evolved and utilized innate defense mechanisms. Even single-celled organisms possess complex heritable defense strategies, some of which are conserved all the way from humans to prokaryotes (*62*), and all multicellular organisms seem to have a sophisticated innate immune system (*63*). However, studies of the innate immune system in invertebrates are scarce and confounded by a small phyletic sampling, primarily limited to flies and nematodes (*64*). Therefore, the evolutionary origin of animal immune cells is still elusive. To investigate this process, we performed an in-depth functional analysis of the antiviral immune response at the cellular level in the early-diverging animal phylum Cnidaria.

Recent scRNAseq studies in the stony coral *S. pistillata* and the sea anemone *N. vectensis* identified distinct immune-related transcriptomic profiles, suggesting potential immune cell functions (*39, 40*). However, these studies lacked functional tests and were conducted under basal conditions, leaving it unclear whether the observed gene expression profiles are functionally linked to immune responses, specific to certain cell types, or indicative of an activatable immune state across various cell types. We filled these critical gaps by demonstrating that immune-responsive cells do exist in *Nematostella* and can be activated by dsRNA viral mimic. Upon activation, these morphologically distinct cells (**Fig. 1** **and** **Fig. 3**) expand rapidly (**Fig. 6A**), launch a robust transcriptomic response (**Fig. 3** and **Fig. 5**), and increase their phagocytic activity (**Fig. 4**).

While we cannot fully negate the possibility that part of the immune cell cluster identified in this study (**Fig. 5 and fig. S5**) represents a cell state rather than a cell type, we provide the following evidence supporting the latter:

1. Although our study was limited to early development, in which cells undergo extensive differentiation, comparison of non-injected control cells belonging to our identified immune cluster with those identified in adults by Cole *et.al* (*40*) demonstrated that 31 out of the 48 reported gene markers were shared between adult immune cells and their embryonic counterparts (**Fig. 1F** and **table S24**). These included all three *Nematostella* RLRs, the transcription factors IRF2, NF-κb1, and ETV6, ISGs such GBP6, GBP3 and OAS1, and members of the Maf family of transcription factors such as MAFG and MAFA. This overlap suggests that the immune cluster we identified in embryos remains stable throughout development. Furthermore, the results by Margolis et al. of the cGAMP treatment on *Nematostella* were obtained by using 4-weeks-old polyps rather than embryos (*42*). The high overlap of differentially expressed genes found in our results suggests that the immune system of embryos and polyps reacts similarly to immune challenges despite the different developmental stages.
2. The putative immune cells described here were morphologically distinct, both at basal conditions and upon activation (**Fig. 1E-G** **and** **Fig. 3** **E and F**).
3. In the immune cell cluster we could not detect the presence of cell markers indicative of other known cell types such as gland cells, nematocytes, neurons and others (**fig. S6A**), further supporting the notion of a cell type rather than a cell state.
4. It is unlikely that the cells belonging to the immune cluster at basal conditions were infected by viruses as our previous study of the *Nematostella* core virome demonstrated that unlike adults that probably acquired viruses via feeding, non-feeding early developmental stages had a negligible viral load that was barely detectable (*65*).
5. In the basal conditions only a few ectodermal cells per embryo were positive for anti-RLRb staining (**Fig. 2, B and C**), making it highly unlikely that those cells are spontaneously infected with a virus.
6. Our pseudotime analysis and the gene modules identified within the immune cluster suggested enhanced proliferation and the activation of the MAPK pathway upon stimulation, pointing to the expansion of a pre-existing cell population that was already present in basal conditions (**Fig. 6F and fig. S8**).

Notably, these two scenarios, cell type versus a cell state, are not mutually exclusive: the presence of specialized immune cells at basal conditions does not exclude the possibility that some of the immune program displaying cells in the poly(I:C) challenged condition are other cell types that responded to the challenge by dramatically changing their transcriptomic profile. The observed shift in cell type composition upon poly(I:C) activation (**Fig. 6A**) might be due to enhanced proliferation of and/or differentiation. Indeed, we found evidence for differentiation and proliferation at the gene expression level (**Fig. 6F** and **fig** **S8C**). Recently, the identification of factors involved in stem cell differentiation in *Nematostella* were described (*66–68*). However, the mechanisms that led to the immune cluster expansion should be further investigated.

Phagocytes, such as macrophages and neutrophils, are thought to be the most evolutionarily ancient blood cells, as they are present in all animals, including simple multicellular organisms like sponges and tunicates (*69, 70*). A recent study suggested that the phagocytic transcriptional program can even be traced back to the protist *Capsaspora owczarzaki* (*71*), a far relative of animals. Interestingly, a cross-species comparison of cell atlases revealed enrichment of shared immune genes in cells of the gastrodermis in *Nematostella*, phagocytes in the planarian *Schmidtea*, and coelomocytes in the nematode *C. elegans* suggesting a shared origin between immunity and digestion (*71*). Even though our study is limited to early developmental stages, we found an overlap of the gastrodermal COA-like-1 marker with the immune cluster we identified (**fig. S6A**). Additionally, we found that cells expressing high levels of RLRb were enriched for peptidase genes (**Fig. 2G**) as previously reported for the gastrodermal immune state in *Nematostella* (*40*). The Rho family GTPases, which some of their members such as RAC and RAS homologs were upregulated upon activation (**Fig. 3J** **and** **Fig. 7F**), are known as widespread regulators of phagocytosis in eukaryotes and seem to have originated from bacteria (*72*). Evidently, activation by dsRNA was accompanied by enrichment of cellular processes potentially indicative of phagocytosis such as plasma membrane organization and actin binding (**fig. S8, C** and **D**) (*73*).

Unlike vertebrates, invertebrates, including insects and nematodes, lack interferons (IFNs) and are believed to primarily depend on RNA interference (RNAi) for antiviral defense (*74*). Here we challenged this dogma by showing that, despite the lack of circulatory system and interferons, the antiviral immune response in *Nematostella* consists of both RNAi and IFN-like components. Such components are also activated in viral infection in vertebrates. For instance, the activation of NF-κB transcription factor binding (**fig. S8D**) (*75*), MAPK signaling pathway activity (**Fig. 3J**) (*76*), and regulation of apoptosis (**Fig. 7F**). The latter was previously described as an essential component of the antiviral immune response in *Nematostella* (*25*). Additionally, other components of the innate immune system seem to be highly conserved. For instance, the complement system which is responsible for rupturing the cell wall of bacteria, promoting phagocytosis, and promoting inflammation in vertebrates, was active in RLRb-high cells (**Fig. 1I**) (*77*). Interestingly, our cross-species analysis revealed similarities in the antiviral and antibacterial immune responses (**Fig. 7**), as it seems that cGAMP, the second messenger cyclic di-nucleotide activator of the antimicrobial cGAS-STING pathway, is capable of inducing genes which are also involved in the antiviral immune response. This was also suggested by Margolis et al. (*42*). However, it is still unknown how these immune responses are activated despite the absence of known cytokines. In contrast to transcription factors and kinases, which are relatively conserved across species, cytokines seem to diverge rapidly between species (*78*). Further research should focus on identifying putative cytokines or other signaling molecules that regulate immune response activation. It is possible that proteinaceous molecules different from canonical vertebral cytokines are involved in this process. In fact, peptidergic paracrine cell signaling was described in Placozoa (*79*). Notably, among the upregulated genes in our immune cluster we identified a variety of G protein coupled receptors (GPCRs) (**fig. S8B**), potentially mediating the immune related signaling pathways. Interestingly, upregulation of hormonal receptors such as the insulin receptor, and enrichment of genes related to sterol binding was also identified (**fig. S8B**), may suggest the involvement of hormone based signaling in this process.

In conclusion, our findings in *N*. *vectensis*, together with our cross-species analysis in the coral *S. pistillata*, which is separated from sea anemones by 500 million years (*80*), suggest that specialized immune cells are ancient and existed in the last common ancestor of class Hexacorallia. Further, by extrapolation, it is highly likely that such cells already existed in the last common ancestor of cnidarian and bilaterian animals, that lived approximately 600 million years ago. This study paves the way for functional identification of factors contributing to the antiviral immune response in cell type specific manner.

## Methods

### Sea anemone culture

*N. vectensis* polyps were grown in 16 ‰ sea salt water (Red Sea, Israel) at 18 °C. Polyps were fed with *Artemia salina* nauplii three times a week. Spawning of gametes and fertilization were performed according to a published protocol (*81*) as follows: in brief, temperature was raised to 25 °C for 9 h and the animals were exposed to strong white light. Three hours after the induction, oocytes were mixed with sperm to allow fertilization.

### Poly(I:C) microinjection

To stimulate the antiviral immune response in *N. vectensis*, we used poly(I:C) (Invivogen, USA) as dsRNA viral mimic. We used 3.125 ng/ml of high molecular weight (HMW) poly(I:C) in 0.9 % NaCl (with an average size of 1.5–8 kb), and 0.9 % NaCl as a control. This concentration was determined as sub-lethal in previous titration experiments (*26*). Transgenic animals were visualized under a SMZ18 stereomicroscope equipped with a DS-Qi2 camera (Nikon). Images were processed using Fiji (*82*). Processing of raw photographs was performed evenly on all files and no manipulations were made to enhance unevenly part of any picture.

### Cell dissociation

Embryos were dissociated into single cells as previously described (*83*). Briefly, individuals were washed twice with calcium/magnesium free and EDTA free artificial seawater (×3 stock solution is: 17 mM Tris-HCl, 165 mM NaCl, 3.3 mM KCl and 9 mM of NaHCO_3_; final solution pH 8.0) and incubated with 50 µg/ml liberaseTM (Roche, Switzerland) at room temperature for 5-10 minutes with occasional gentle pipetting, until fully dissociated. The reaction was stopped by adding 1/10 volume of 0.5 M EDTA solution. The suspension was filtered using a 35 µm cell strainer. Cells were then centrifuged at 500 × *g* at 4 °C and resuspended in 1× calcium/magnesium free sterile PBS (Hylabs, Israel) supplemented with 2% BSA. Cells were counted on hemocytometer and viability was determined using trypan blue (Thermo Fisher Scientific, USA).

### Flow cytometry

FACSAria III (BD Biosciences, USA) equipped with 405 nm, 408 nm, 561 nm and 633 nm lasers was used to quantitively assess phagocytosis, mCherry expression, and RLRb expression. Per run, 30,000 events were recorded. In experiments involving the *RLRb::mCherry* transgenic reporter line Helix NP (SYTOX Blue) (Ex430/Em470) (Biologend, USA) and calcein AM were used to determine viability. For immunostaining experiments, the nuclear far-red marker DRAQ5 (Abcam, United Kingdom) was used to identify intact fixed and permeabilized cells. In phagocytosis experiments, Zombie NIR (Biolegend) and Calcein violet AM were used to assess viability and to discern cells from phagocytosis probes (fixed bacteria or beads). FCS files were further analyzed using FlowJo V10 (BD Biosciences). Each measurement consisted of 3 biological replicates, unless indicated otherwise.

### FACS isolation of live *RLRb::mCherry* cells

400-500 24 hpi planulae were used for cell dissociation. Cell dissociation was carried out as described in the previous section. Dissociated cells were resuspended in 5mL polypropylene Flacon tubes (BD Falcon 352063) with sterile 1× PBS (calcium/magnesium free) (Hylabs) buffer containing 2 % BSA and the viability dyes: 2 µg/mL calcein AM (Enzo) and 100 nM sytox blue (BioLegend). Before sorting, FACSAria III (BD Biosciences) was sterilized by running bleach for 5 minutes followed by 5 minutes of 70 % ethanol at maximal speed. 100 µm nozzle was used for sorting. Immediately before sorting, cells were passed once more through a 35 µm cell strainer and mixed vigorously by pipetting. Cells were sorted at low pressure at a speed of up to 3,000 events/sec. The sample tubes were kept at 4 °C throughout the process. To maximize RNA yield by reducing cell adherence to the sides of the tubes, 15 mL conical bottom tubes were filled with PBS with 20 % fetal bovine serum (FBS) (Gibco) and incubated overnight at 4 °C degrees. Immediately before sorting, the tubes were emptied and filled with 1x cold PBS (calcium/magnesium free) (Hylabs). Collection tubes were kept at 4 °C during the sorting process. 150,000-600,000 viable cells were collected per cell fraction. Once completed, cells were centrifuged at 1000 × *g* at 4 °C for 10 minutes, resuspended in 1 mL TRIzol Reagent (Thermo Fisher Scientific), and immediately processed for RNA extraction (see next section).

### RNA extraction

Extraction of RNA from pelleted FACS sorted cells was conducted according to the TRIzol Reagent (Thermo Fisher Scientific) manufacturer’s instructions with slight modifications. Briefly, cells were lysed in TRIzol Reagent by vigorous pipetting. At the precipitation stage, 1:1 volume of isopropanol was added to the aqueous phase. 2 µL of glycogen (Thermo Fisher Scientific) were added into each tube. The solution was then mixed by inverting the tubes 10 times. Samples were then incubated at -20 °C for 30 minutes and centrifuged at maximal speed (∼21,000 x *g*) in 4 °C microcentrifuge. The pellet was then washed twice with 75 % ethanol in DEPC-treated water and eluted in 15 µL of nuclease free water. RNA was quantified using Qubit RNA High Sensitivity (HS) Kit (Thermo Fisher Scientific). RNA integrity number was obtained using Bioanalyzer RNA 6000 Pico Kit (Agilent).

### RNAseq library construction and sequencing

Only samples with RNA integrity number >7.5 were processed. Libraries were constructed from 60 ng of total RNA from mCherry positive and negative cells of the *RLRb::mCherry* transgenic line 24 hours after poly(I:C) injection. For intracellular immunofluorescent staining experiments, a total of 100 ng was used as input. RNAseq libraries were generated using NEBNext Ultra II Directional RNA Library Prep with Sample Purification Beads (New England Biolabs, USA) following the manufacturer’s protocol. Samples were PCR amplified and indexed using NEBNext Multiplex Oligos for Illumina Kit (New England Biolabs) using 14 cycles. PCR amplified library products were then quality checked using High Sensitivity D1000 DNA Screen Tape assay (Agilent) loaded into the Agilent 2200 TapeStation system (Agilent). Samples showing a narrow distribution with a peak size approximately 300 bp were then sequenced on NextSeq 2000 (Illumina, San Diego, CA) using dual index mode with single-end read length of 100 bp.

### Phagocytosis assays

*RLRb::mCherry* zygotes were injected with Poly(I:C) or NaCl and were incubated for 24 hours at 22°C. In parallel, wild type and *RLRb::mCherry* uninjected controls were kept at the same conditions. Cells were then dissociated as described in *Cell dissociation* section. Cells were counted on a hemocytometer using trypan blue to assess viability. Phagocytosis was examined as previously described (*37*). Briefly, 70,000 live cells were seeded per well in a 96-well plate in L-15 based medium supplemented with 2 % heat inactivated fetal bovine serum (FBS), 20 mM HEPES, and calcium and magnesium free 10 × PBS brought to 1.42 X PBS molarity. 0.05 % of NaN_3_ was added to reduce contamination. Cells were then incubated for 16 hours at 18 °C with either of the phagocytic probes: 15 µg/mL *Staphylococcus aureus* (pHrodo™ Green *S. aureus* Bioparticles Conjugate for Phagocytosis; Thermo Fisher Scientific), 15 µg/mL *Escherichia coli* (pHrodo Green *E. coli* BioParticles Conjugate for Phagocytosis; Thermo Fisher Scientific), 15 µg/mL DQ ovalbumin (Thermo Fisher Scientific), or none. Flow cytometry analysis was carried out to determine whether mCherry positive cells were enriched for phagocytosis. Dead cells, debris and leftover phagocytic probes were excluded by gating calcein violet AM positive cells. In addition, Zombie NIR was used to gate viable cells. Spillover of the green phagocytic probe into the mCherry channel was accounted for by gating based on WT cells that lack a signal at the mCherry channel (**fig. S1A**). The percentage of cells positive to the phagocytic probe in the fraction of mCherry positive and mCherry negative cells was determined using FlowJO V10 (BD Biosciences) and plotted using ggplot2 (*84*) and Hmisc (*85*) R packages. Welch two sample two-sided t test was conducted to determine statistical significance.

### Whole animal immunofluorescent staining

Immunostaining was performed according to a previously described protocol (*86*), using a custom made polyclonal antibody against RLRb (GenScript) which was previously characterized for western blot (*26*). In brief, about 150 embryos 24 hpf were fixed for 1 hour in 4 % paraformaldehyde with 0.1% Tween-20 in PBS (1.86 mM NaH_2_PO_4_, 8.4 mM Na_2_HPO_4_, 175 mM NaCl, pH 7.4). Following fixation, the samples were washed with 0.1 % Tween-20 in PBS and stored in absolute methanol at –20 °C until further use. After gradual rehydration, the samples were blocked for 2 hours in a solution containing 20 % sheep serum (Sigma Merck Millipore, USA) and 3 % bovine serum albumin (BSA, fraction V) in PBS with 0.1% Triton X-100. Primary antibodies against RLRb (GenScript) and against mCherry (ChromoTek, Germany) were used at 1:1000 and 1:250, respectively. After incubating overnight at 4°C, the samples were washed five times with PBS containing 0.1% Triton X-100, re-blocked, and subsequently incubated with the secondary antibodies: anti-rabbit conjugated to Alexa fluor 488 (Jackson’s Immunoassays), or anti-rat Alexa fluor 568 (Jackson’s Immunoassays), for RLRb and mCherry respectively. The samples were washed again five times with PBS containing 0.1 % Triton X-100 and mounted on microscope slides using VectaShield (Reactolab, Switzerland). Specimens were visualized using Eclipse Ni-U microscope equipped with a DS-Ri2 camera and an Elements BR software (Nikon, Japan) or with FV1200 confocal microscope (Olympus, Japan). Images were processed using Fiji (*82*). Processing of raw photographs was performed evenly on all files and no manipulations were made to enhance unevenly part of any picture.

### Intracellular immunofluorescent staining and FACS

Cells derived from 48 hours old planulae were dissociated as described above in the Cell dissociation section. Dissociated cells were then washed twice with PBS. Cells were fixed with 4% Paraformaldehyde (PFA) in PTW (PBS, 0.1 % Tween-20) for 30 minutes on a rotator at 4 °C. In the following steps centrifugation was performed for 3 minutes at 3000 x *g* at 4 °C. All subsequent steps were performed on ice with the addition of murine RNAse inhibitor (New England Biolabs) at a concentration of 1 unit/µL to prevent RNA degradation. Cells were resuspended in Wash Buffer (PBS, 0.2%BSA, 0.1% Triton-x), washed twice, and counted using 1:10 concentration of DAPI (Thermo Fisher Scientific) on a hemocytometer. 5 million DAPI positive cells were used per 1 mL Staining Buffer (PBS, 2% BSA, 0.1% Triton-X). Staining with primary antibody, anti-RLRb rabbit (GenScript) or anti-IgG rabbit (Sigma), at 1:1000 concentration was carried for 1 hour at 4 °C on a rotator. This concentration was determined based on titration experiments that demonstrated a decent signal to noise ratio. Cells were then washed twice in Wash Buffer. Cells were then stained with 1:1000 secondary antibody, anti-rabbit Alexa fluor 488 (Jackson’s Immunoassays). Cells were then washed twice in Wash Buffer and resuspended in Sort Buffer (PBS, 0.5 % BSA) containing 1:1000 DRAQ5 (Abcam). Emitted fluorescence was acquired at the Alexa fluor 488 channel for RLRb and APC-Cy7 channel for DRAQ5. Gates were set with reference to the negative anti-IgG control. The sorting speed was adjusted to ensure sorting efficiency above 80 %. Cells were sorted into BSA pre-coated 15mL tubes as described in previous sections. About 0.5 million cells were collected for the RLRb high cells and about 2 million cells for the RLRb low cells per biological replicate. The experiment was repeated for a total of 4 biological replicates.

### RNA extraction from paraformaldehyde fixed cells

PFA fixed cells following FACS were centrifuged at 2600 × *g* for 5 minutes at 4 °C. At this step the pellet was invisible. The supernatant containing FACS buffer was carefully removed using a pipette. For reverse crosslinking, cells were incubated in 300 µL PBS containing 2.5 µL 20mg/mL Proteinase K (Thermo Fisher Scientific) and 3 µL 10 % SDS at 50 °C for 3 hours. The lysate was then kept at -80 °C until further processing. To extract RNA, TRIzol-LS Reagent (Thermo Fisher Scientific) was used according to the manufacturer’s instructions with the modifications mentioned in the RNA extraction section above.

### Imaging Flow Cytometry

Intracellular immune-stained cells labeled with anti-RLRb or anti-IgG control antibodies were assessed quantitatively using an Amnis ImageStreamX Mk II apparatus (Luminex, USA) equipped with 405 nm, 488 nm, 642 nm and 785 nm (SSC) lasers and 6 acquisition channels, using a 60× magnification objective, with low flow rate/ high sensitivity using INSPIRE software (Luminex, USA) for data acquisition. The INSPIRE software was set up using the following parameters: Channel 01 (bright field), Channel 02 for detecting laser 488nm, and Channel 05 detecting laser 642nm. Cells were resuspended in 100 μL of PBS in 1.5 ml

Eppendorf tubes. About 10^7^ cells per sample were used for data acquisition. A total of 10,000 single events were acquired within the focused singlet gate. Focused events were determined by the Gradient RMS parameter for Ch01 and single cells were determined by plotting the area of the events (X-Axis) vs. aspect ratio (Y-Axis) for Ch01. All subsequent analysis was done on this population of cells. Image analysis was run using IDEAS 6.3 software (Luminex), circularity feature was determined using the Shape Change Wizard, which provided circularity score for each population by measuring the degree of the mask’s deviation from a circle. Downstream analysis was performed using R 4.4.1 version. For area, circularity, granularity, and Alexa fluor 488 intensity data obtained by previous analyses using IDEAS was directly plotted using ggplot2 (*84*) and Hmisc (*85*) R packages. To determine statistical significance, Welch two sample two-sided t test was conducted. For PCA, raw data was loaded into R and the three biological replicates were merged (82,340 observations, 104 parameters), events were split into the top 15 % against the rest of the data based on Alexa fluor 488 intensity, within the RLRb stained samples. Zero variance columns (parameters) were removed, and NAs were omitted. Alexa fluor 488 intensity was then removed from the numeric matrix. The resulting numeric data (28,409 observations of 91 parameters) was standardized using the base R *scale* function. PCA was computed using the *prcomp* function, individual data points were visualized as centroids representing the mean of PC1 and PC2 for each replicate. The result was visualized using ggplot2 (*84*).

### RLRb immunoprecipitation

100 µl of magnetic beads (SureBeads™ Protein A Magnetic Beads, Bio-Rad, USA) were washed in 1 ml of 1×PBS for 5 times. 5 µg of RLRb antibody (Genescript, USA) or total rabbit IgG was added to the washed beads with 1.3 ml of 1×PBS in triplicates and left rotating overnight at 4°C. Animals that correspond to a volume of 100 µl were frozen in liquid nitrogen and lysed (with homogenizer) using 1 ml of the following lysis buffer: 25 mM Tris-HCl (pH 7.4), 150 mM KCl, 25 mM EDTA, 0.5% NP-40, 1 mM DTT, Protease inhibitor cOmplete ULTRA tablets (Roche) and Protease Inhibitor Cocktail Set III, EDTA-Free (Merck Millipore, USA). The DTT, Protease inhibitor and RNAse inhibitor were added fresh just before use. After 2 h rotation in 4°C the samples were centrifuged at 16000 × *g*, 15 min, 4 °C and supernatant was collected. Next, the lysate was precleared as following: 100 µl of magnetic beads were washed in 1 ml of 1×PBS for 3 times and 1 time in lysis buffer and the lysate was added to the washed beads. Lysis buffer with RNAse inhibitor was added to make up 1.2 ml and the samples were incubated at 4 °C rotation for one hour. Next, the pre-cleared lysate was collected and added to the antibody-bound beads (that were pre-incubated with the antibody overnight). These samples were incubated for 2 h in rotation at 4 °C. After incubation the lysate was removed, and the beads were washed 5 times with the following wash buffer: 50 mM Tris-HCl (pH 7.4), 300 mM NaCl, 5 mM MgCl_2_, 0.05 % NP-40, Protease inhibitor cOmplete ULTRA tablets (Roche, Switzerland) and Protease Inhibitor Cocktail Set III, EDTA-Free (Merck Millipore).

### Semi-quantitative LC-MS/MS analysis

After the last step of immunoprecipitation, the beads were washed twice with 25 mM Tris-HCl pH 8.0. The packed beads were re-suspended in 100 μl 8M urea, 10 mM DTT, 25 mM Tris-HCl pH 8.0 and incubated for 30 min at 22 °C. Next, Iodoacetamide (55 mM) was added, and beads were incubated for 30 min (22°C, in the dark), followed by addition of DTT (20 mM). The Urea was diluted by the addition of 6 volumes of 25 mM Tris-HCl pH 8.0. Trypsin was added (0.3 μg per sample) and the beads were incubated overnight at 37°C with gentle agitation. The beads were spun down and the peptides were desalted on C18 home-made Stage tips. Two-thirds of the eluted peptides were used for MS analysis.

### nanoLC-MS/MS analysis

MS analysis was performed using a Q Exactive Plus mass spectrometer (Thermo Fisher Scientific) coupled on-line to a nanoflow UHPLC instrument, Ultimate 3000 Dionex (Thermo Fisher Scientific). Peptides were separated over a 60 min gradient run at a flow rate of 0.3 μl/min on a reverse phase 25-cm-long C18 column (75 μm ID, 2 μm, 100 Å, Thermo PepMapRSLC). The survey scans (380–2,000 m/z, target value 3E6 charges, maximum ion injection times 50 ms) were acquired and followed by higher energy collisional dissociation (HCD) based fragmentation (normalized collision energy 25). A resolution of 70,000 was used for survey scans and up to 15 dynamically chosen most abundant precursor ions, with “peptide preferable” profile were fragmented (isolation window 1.6 m/z). The MS/MS scans were acquired at a resolution of 17,500 (target value 1E5 charges, maximum ion injection times 120 ms). Dynamic exclusion was 60 sec. Data were acquired using Xcalibur software (Thermo Scientific). To avoid a carryover, the column was washed with 80 % acetonitrile, 0.1 % formic acid for 25 min between samples.

### MS data analysis

Mass spectra data were processed using the MaxQuant computational platform, version 1.5.3.1254. Peak lists were searched against translated coding sequences of *Nematostella* gene models. The search included cysteine carbamidomethylation as a fixed modification and oxidation of methionine as variable modifications and allowed up to two miscleavages. The match-between-runs option was used. Peptides with a length of at least seven amino-acids were considered and the required FDR was set to 1% at the peptide and protein level. Protein identification required at least 2 unique or razor peptides per protein. Relative protein quantification in MaxQuant was performed using the label-free quantification (LFQ) algorithm (*87*). LFQ in MaxQuant uses only common peptides for pair-wise ratio determination for each protein and calculates a median ratio to protect against outliers. It then determines all pair-wise protein ratios and requires a minimal number of two peptide ratios for a given protein ratio to be considered valid. Statistical analysis (n=3) was performed using the Perseus statistical package (*88*). Only sample groups with at least 2 valid values were used. Protein contaminants and proteins identified by less than 2 peptides were excluded from the analysis. The procedure described above was carried out on three technical replicates for each RLRb-IP.

### Bulk RNAseq analysis and GO functional analysis

The quality of raw reads was evaluated and visualized using FastQC (*89*), with summary reports generated through multiqc (*90*). Reads were trimmed using Trimmomatic (*91*) with the following parameters: ILLUMINACLIP:TruSeq3-SE:2:30:10, LEADING:3, TRAILING:3, SLIDINGWINDOW:4:15, and MINLEN:36. The trimmed reads were aligned to the Darwin Tree of Life (DToL) *N. vectensis* reference genome jaNemVect1.1 (*92*) using STAR (version 2.7.10a) (*93*). Gene-level read counts were summarized using featureCounts (version 2.0.1) (*94*). Genes with >= 10 across all samples were kept. Differentially expressed genes (DEGs) across different cell fractions were identified using DESeq2 (*95*) with the design = ∼ condition argument. DEGs were defined as genes with an absolute log2 fold change > 1 and an adjusted p-value < 0.05. Principal component analysis (PCA) was performed on variance-stabilized transformed (vst) data using the DESeq2 vst function with the blind = FALSE argument. Data visualization was carried out using the Bioconductor EnhancedVolcano R package (*96*) for volcano plots and ggplot2 (*84*). For comparison, the data published Lewandowska *et.a* (*26*)*l* of poly(I:C) versus NaCl injected *Nematostella* zygotes at 24 hpi was downloaded using SRA Toolkit (*97*). To address the differences between gene models derived from two versions of the

*Nematostella* genome, we constructed a correspondence dictionary (**table S25**). This was achieved through reciprocal BLASTp v2.13.0 (*98*) analysis, where each version was alternately used as the database and query, ensuring accurate mapping between the gene models of both genome versions. The same pipeline was applied. Venn diagrams were generated using the VennDiagram R package (*99*). Functional annotation was performed as previously described (*25*) using the DtoL gene models. Gene set enrichment analysis (GSEA) and over-representation analysis (ORA) were conducted using clusterProfiler (*100*). We utilized the QuickGO (*101*) database to retrieve GO annotations. Using QuickGO’s query tools, we obtained a comprehensive mapping of gene identifiers to their associated GO terms, ensuring coverage across biological processes, molecular functions, and cellular components. The resulting GO annotation file was curated and formatted to align with the input requirements for the clusterProfiler R package (*100*).

### ACME dissociation for 10X scRNAseq

Embryos of *N. vectensis* from a transgenic line expressing *RLRb::mCherry* were treated as described above, including a control group of non-injected embryos, a control group injected with NaCl, and a group injected with poly(I:C). The experiment was conducted in two biological replicates, with 400 embryos at the gastrula stage (24 hpf) collected from each treatment group. Embryos were dissociated as described in the Cell dissociation section above. Cell viability was assessed using Calcein-AM green and Helix NP as before, and the activation of the RLRb promoter was evaluated by measuring mCherry fluorescence. Both measurements were performed using flow cytometry. After viability and mCherry assessment, cells were fixed in ACME solution (Ca-free *N. vectensis* medium, glacial acetic acid, glycerol, methanol, and EDTA in a 13:2:2:3:1 ratio) for 30 minutes at room temperature on a rotary platform (*44*). Cells were then washed twice in resuspension buffer (RB1: PBS 1× nuclease-free, sorbitol 0.8 M, 40 U/ml RNasin Ribonuclease Inhibitor), via centrifugation at 1,000 × *g* for 5 minutes at 4 °C. Cells were counted using DAPI staining (9 µl of cell suspension mixed with 1 µl DAPI, 1 mg/ml). Cell concentration was calculated and adjusted to reach 400,000 cells in 100 µL for each aliquot.

### ClickTag Barcoding for scRNAseq

Fixed cells were barcoded using a modified version of the ClickTag protocol (*52*). To optimize the labeling reaction in ACME fixative, the amine-reactive cross-linker TCO-NHS used by Gehring et al. (*52*) was replaced with TCO-PEG4-TFP (Click Chemistry Tools, China), which offers improved stability against hydrolysis in aqueous media. Barcoding DNA oligonucleotides (ClickTags) with a 5’-amino modifier (Integrated DNA Technologies, USA) were activated by derivatization with methyltetrazine-NHS ester (MTZ-NHS) as originally described. For cell tagging, each treatment sample was labeled using a combination of three distinct MTZ-derivatized oligonucleotides.

### ClickTag Labeling

For ClickTag labeling, cells were pre-incubated with 4.5 µL of 1 mM TFP-TCO for 5 minutes at room temperature in the dark. Cells were then labeled with a mixture of three MTZ-activated tags (4 µL each), specific to each treatment. After 30 minutes of incubation on a rotary platform in the dark, the reaction was quenched with 13 µL of 100 mM Tris-HCl and 0.65 µL of 10 mM MTZ-DBCO (*52*). Labeled cells from each treatment were pooled by biological replicate (1.2 million cells per pool). Pools were washed twice with 1 mL of resuspension buffer (RB2; 1× PBS, 0.5% BSA, and 40 U/mL RNase Inhibitor, Murine, New England Biolabs) by centrifugation at 1,500 × *g* for 10 minutes at 4 °C. Cells were resuspended in RB2 containing 10 % DMSO and stored at -80 °C until further processing.

### Cell Sorting and Single-Cell RNAseq

Single-cell transcriptomes were obtained using the Chromium Single Cell 3’ Gene Expression kit v3.1 (10x Genomics). Frozen samples were thawed on ice, and cells were collected by centrifugation at 2000 × *g* for 5 minutes at 4 °C. After one wash with 2 mL of Resuspension Buffer 2 (RB2; 1 × PBS nuclease-free, 0.5% BSA, 40 U/ml RNase Inhibitor, Murine, New England Biolab), cells were pelleted again and resuspended in 1 mL of RB2. Cells were stained with 1:1000 DAPI for nuclei staining. For single-cell isolation, 15,000 cells from each biological replicate, were sorted into two well of a 96-well plate using a FACSAria II SORP cell sorter, following 10x Genomics’ guidelines. Non-cellular particles were excluded by selecting only DAPI-positive cells, and doublets/multiplets were removed using forward scatter width (FSC-W) versus forward scatter height (FSC-H). Cells were immediately encapsulated after sorting, and barcoded cDNA and sequencing libraries were prepared according to 10x Genomics’ protocols, each biological replicate in a separate lane. For ClickTag library preparation, ClickTag cDNA was separated from cellular cDNA after the cDNA amplification step, using differential size-selection purification with AMPure XP beads (Beckman Coulter). ClickTag sequencing libraries were prepared as described previously (*52*). The size distribution and concentration of the final libraries were assessed using a TapeStation (Agilent) and Qubit (Thermo Fisher Scientific). Libraries were sequenced on an Illumina NextSeq 500 sequencer using high-output 75-cycle V2 kits (Illumina).

### Transcriptome mapping and Clicktag demultiplexing

We used CellRanger 6.1.1 (10X Genomics) to map reads and count unique molecular identifiers (UMIs) per gene per cell, using *N. vectensis* DToL genome assembly and gene models (*92*). We also added the mCherry sequence and gene model to this reference genome, in order to be able to quantify the expression of the *RLR::mCherry* reporter. We used the --force-cells flat to set a constant number of 20,000 per experiment and we used whole gene bodies to guide the mapping of reads to the genome (--transcriptome flag). We discarded all cell barcodes with less than 1,000 UMIs/cell, upon examination of the distribution of UMIs per cell. The scRNAseq UMI matrices of all the *bona fide* cells were then converted to MetaCell 0.37 format (*53*) for further processing in R.

We then used Clicktag barcodes to first remove potential doublets, removing all cells that failed any of the following criteria: (1) first, we only kept cells with a Clicktag count >20 UMIs. Clicktag counts were determined as described by Chari *et al.* (*102*): we mapped the reads to the Clicktag barcodes (8 bp barcodes + constant CAG sequences at the end) using *kallisto* 0.46.2 (*103*), specifying the 10x v3 chemistry and tolerating one substitution per barcode; and used *bustools* 0.41.0 (*104*) to correct, sort and count the reads per cell, and obtain a final Clicktag UMI matrix. (2) second, we compared the number of Clicktag counts for the most abundant barcode to the third most abundant barcode for each cell (given that we had used two barcodes per experimental batch, the abundance of the third most abundant barcode should be lower than the first and second ones, and correspond to a different batch of cells); and we kept cells where the first-to-third ratio of normalized Clicktag UMI counts was > 1.5. Likewise, we flagged as possible doublets all cells where the first and second most abundant Clicktag barcodes corresponded to different batches. Cells failing these criteria were flagged as potential doublets, and further classified as intra- or inter-species doublets depending on whether the two most abundant Clicktag barcodes corresponded to barcode pairs from the same/different species, respectively. (3) Third, we used Seurat 4.1.1 (*105*) to produce Louvain clusters of cells based on their Clicktag count matrix, and removed all cells that fell within clusters that contained a high fraction (>70%) of cells flagged as potential doublets according to either of the previous criteria. These Seurat clusters were generated as follows: we used scaled and log-normalized counts with a scaling factor of 10,000 (NormalizeData function); selected variable features using default parameters (FindVariableFeatures function); ran a PCA and identified the 50 nearest neighbors of each cell (RunPCA and FindNeighbors functions with k = 50); and used this distance matrix to identify Louvain clusters (FindClusters function, with resolution = 1).We finally classified all cells passing the filters described above (11,550 cells) into each of the experimental conditions (Control, NaCl control, and pIC-treated) using their most abundant Clicktag barcode combinations.

### Metacell clustering and analysis

We used Metacell 0.37 (*53*) to select gene features and construct cell clusters (termed metacells). We selected feature genes using normalized size correlation (*53, 106*) threshold of -0.1 and normalized niche score (*53*) threshold of 0.01, additionally filtering for genes with > 1 UMI in at least three cells and a total gene UMI count > 30 molecules (mcell_gset_iter_multi function in Metacell), selecting a total of 915 variable genes for clustering. For K-nearest neighbours graph building we used K = 100 target number of edges per cell (mcell_add_cgraph_from_mat_bknn function), and for metacell construction we used K = 30, minimum module size of 10, and 1,000 iterations of bootstrapping with resampling 75% of the cells, and threshold α = 2 to filter edges by their co-clustering weight (mcell_coclust_from_graph_resamp and mcell_mc_from_coclust_balanced functions). This way we obtained an estimate of co-clustering frequency between all pairs of single cells and identified robust clusters of single or grouped metacells. For downstream analyses, we represent gene expression by computing a regularized geometric mean within each metacell (or cell type, where appropriate) and dividing this value by the median across metacells, as implemented in Metacell. We refer to these normalized gene expression values as fold change (FC) across the manuscript.

To identify cell types in our scRNA-seq dataset, we compare our metacells with a wild-type gastrula scRNA-seq atlas [ref Cole], processing this reference atlas as described above. For comparison, we selected variable genes (max FC > 2 in our dataset) and used them to calculate Pearson correlation coefficients between metacells. The resulting comparative heatmap (fig. S7) was used to manually transfer cell type annotations.

### Gene module analysis

We used the metacell normalized gene expression fold change (FC) to obtain gene modules using the *WGCNA* algorithm (*107*). First, we selected variable genes with a FC > 1.5 in at least one metacell. Second, we calculated the gene co-expression matrix by calculating the Pearson correlation coefficient of each gene based on their metacell fold changes, and using the average hierarchical clustering algorithm and a soft power parameter=7 (determined independently for each sample using the WGCNA *pickSoftThreshold* function). Third, we used the hierarchical clustering dendrogram to define gene modules using the *cutreeHybrid* function in the *dynamicTreeCut* R library, maximizing granularity with a split parameter=4; assigned each gene to one module with a correlation threshold R 0.7; and we calculated the module Eigen vectors of each of the resulting modules in each metacell, using the *moduleEigengenes WGCNA* function.

### scRNAseq clustering and analysis

The CellRanger (*108*) generated count matrix described above, along with cell barcodes, gene annotations, condition (i.e. poly(I:C), NaCl, ctrl), and metacell identifiers metadata were loaded into R using the *Seurat* (*109*) functions CreateSeuratObject and AddMetaData. For pre-processing, number of genes per cell and number of transcripts were visualized. The following filtering was applied: nFeature_RNA > 500 & nFeature_RNA < 3000 & nCount_RNA < 10000. In addition, only genes expressed in at least 10 cells were kept. This resulted in 16,535 genes and 11,065 cells. Next, the sctransform R package (*110*) was used for normalization and variance stabilization using the SCTransform function with default parameters. Data reduction was performed using PCA. For clustering, dimensions 1 to 30 were used with resolution = 0.2. These parameters allowed clear discrimination of previously reported marker genes (*40*), including gland cells and nematocytes, resulting in 8 different clusters. The two biological replicates, corresponding to Nvec01 and Nvec02 prefix, were visualized on a UMAP projection (supplementary) revealing a similar distribution (supplementary). Therefore, for downstream analysis the two replicates were merged. We determined mCherry expression per condition using the SCT transformed matrix and visualized the results using ggplot2 (*84*). Correlation analysis between RLRb (Nvec_vc1.1_XM_048731786.1) and mCherry (mCherry_plus_strand) was done by plotting UMI fractions for each gene per metacell using the pearson correlation method in ggpubr (*111*). To determine in which cell population the genes that were upregulated in the mCherry positive cells are enriched, we used the Seurat AddModuleScore and visualized the aggregated score on the umap plot. For differential expression analysis within different conditions and different clusters we used the MAST (*112*) method with default parameters.

### Sub-clustering and Pseudotime analysis

To gain a higher resolution into gene expression changes within cluster 1, we re-clustered its data by subsetting it and performing clustering as described in the previous section. In this case the data was normalized and scaled using NormalizeData, FindVariableFeatures and ScaleData in Seurat (*109*). Clustering was done using PC1 to PC15 using resolution = 0.2. To identify sub-clusters marker genes the FindAllMarkers function was used with the following arguments: min.pct = 0.25 and logfc.threshold = 0.25. Top markers were selected based on their average log2 fold change and visualized using a dotplot with ggplot2 (*84*). Pseudotime analysis was conducted on the re-clustered subset of the data using Slingshot (*55*). The data was first converted to a SingleCellExperiment object using the Bioconductor SingleCellExperiment package (*113*). The *slingshot* function was applied to this object and pseudotime was fetched using slingPseudotime on columns corresponding to “Lineage1”. slingCurves was then applied to fetch principal curve in PCA space. embedCurves was ran on the slingshot object with the arguments: ’UMAP’, smoother = ’loess’, span = 0.1, followed by slingCurves. The UMAP curve was ordered and visualized using ggplot2. To model gene expression as a function of pseudotime generalized additive models (GAM) (*114*) was used. Observed expression values along with a GAM-fitted curve were visualized using ggplot2 for genes of interest.

### Cross species analysis

To assess the conservation of transcriptional responses between *N. vectensis* and *S. pistillata*, we integrated species-specific differential expression (DE) results with orthology relationships inferred using OrthoFinder v2.4.0 (*115*), a tool that combines BLAST-based similarity searches and phylogenetics for high-accuracy orthogroup and ortholog identification. The orthogroup output was processed to generate a one-to-one mapping of individual *Nematostella* and *Stylophora* gene pairs by expanding multi-gene entries and standardizing gene identifiers. This allowed consistent merging of orthology assignments with DE results. Differentially expressed genes of S. *pistillata* treated with 2’3’-cGAMP published by Li *et.al* (*116*) were used for comparison against the MAST differential expression analysis of cluster 1 against the all other clusters conducted in the current study. Differentially expressed genes were identified independently for each species, and significance was defined as an adjusted p-value (padj) < 0.05. DE results were then joined with the orthogroup-expanded table, and for each *Nematostella* gene, all orthogroups containing it were evaluated. Genes were classified into one of three mutually exclusive expression categories using the following prioritization scheme:

1. Conserved - the *Nematostella* gene is part of at least one orthogroup in which both orthologs are significantly differentially expressed in the same direction (i.e., log₂ fold changes share the same sign).
2. Divergent - the gene is not in any conserved orthogroup, but is part of at least one orthogroup where both species show significant differential expression in opposite directions.
3. *Nematostella*-specific - the gene is significantly DE in *Nematostella*, but either has no ortholog with DE in *Stylophora* or is not included in any orthogroup.

This classification was applied at the *Nematostella* gene level to ensure each gene was assigned to a single biologically relevant category. Among significantly DE *Nematostella* genes, those without any orthogroup assignment were identified as orphan DE genes, representing candidate lineage-specific or taxonomically restricted genes. This gene-level categorization allowed us to distinguish transcriptional responses that are evolutionarily conserved, functionally divergent, or potentially unique to *Nematostella*. The resulting groups were used for downstream analysis and visualization of expression correlation, Venn diagrams, and functional enrichment.

## Acknowledgements

The authors would like to thank Ms. Adi Turjeman and Dr. Michal Bronstein of the Genomic Technologies Center of the Hebrew University of Jerusalem for their help with RNA sequencing, Dr. Naomi Melamed-Book of the imaging facilities at the Hebrew University for her help with confocal microscopy, and Dr. Tzachi Hagai (Tel Aviv University) and Prof. Oren Ram (The Hebrew University) for helpful discussions about the analysis and interpretation of the results. This work was supported by European Research Council Consolidator Grant 863809 AntiViralEvo to Y.M. B.R. was supported by European Research Council (ERC) Starting Grant 948476 and by the NSF-BSF Integrative and Organismal Systems (IOS) grant (BSF grant no. 2019647, NSF grant no. 1951826). Research in A.S-P. lab was supported by Research in A.S-P. group was supported by the European Research Council Starting Grant 851647

## Data availability

All sequencing data obtained in this study are available at SRA BioProject accession number PRJNA1207058 and under GEO accession number GSE284212. Genome and transcriptome annotation files are available at https://doi.org/10.6084/m9.figshare.28265126. Supplementary tables are available at https://doi.org/10.6084/m9.figshare.28266230. MS/MS raw files, as well as results of MaxQuant analysis were deposited to the ProteomeXchange Consortium via the PRIDE55 partner repository with the dataset identifier: PXD064383.

## List of Supplementary Figures and Tables

**fig. S1. Gating strategies of intracellularly immunostained cells. (A)** ImageStream gating strategy. Focused cells were gated based high gradient RMS, aggregates and debris where excluded based on aspect ratio and manual inspection of individual cell images, intact cells were selected by the nuclear marker DRAQ5, RLRb signal was defined relative to IgG control. **(B)** Gating strategy for flow cytometry.

**fig. S2.** **Validation of intracellular immunostaining for the detection of RLRb in single cells (A)** Titration of the RLRb antibody relative to IgG control. **(B)** flow cytometry results of IgG stained control, **(C)** NaCl control and **(D)** poly(I:C) injected cells. **(E)** flow cytometry results in WT derived cells. **(F)** Flow cytometry results in RLRb CRISPR knockout derived cells.

**fig. S3.** Gating strategy for experiments utilizing the *RLRb:mCherry* transgenic reporter line. (A) Single cells were gated as shown and viability was determined using helix-NP and CalceinAM green.

**fig. S4.** **Gating strategy for phagocytosis experiments. (A)** Single cells were gated as before. Calcein violet AM and Zombie NIR were used to determine viability. mCherry was plotted against the phagocytic probe and gates were defined with respect to WT untreated controls.

**fig. S5.** Metacell and scRNAseq are reproducible and reveal poly(I:C) specific cell states. (A) Top panel shows the distribution of metacells across two biological replicates. Bottom panel shows the distribution of metacells across conditions. The heatmap shows gene expression (rows) across metacells (columns). (B) scRNAseq distribution of cells across two biological replicates visualized on a UMAP projection. (C) Scaled RLRb expression is shown in green. Scaled mcherry expression is shown in red. Cells co-expressing both are shown in yellow.

**fig. S6. Cell type specific markers. (A)** Expression of cell type specific markers reported by Cole *et.al* (***40***) is shown on UMAP projections. The marker and the cell type or tissue of origin are indicated.

**fig. S7. Identification of cell-type specific gene modules. (A)** WGCNA analysis revealed 19 distinct gene modules (named GS) across metacells. The heatmap shows the enrichment of the modules across metacells. **(B)** Average expression score per module was computed across cell clusters and visualized as violin plots.

**fig. S8. ORA of immune specific gene modules. (A)** ORA of GS17, **(B)** GS16, **(C)** GS14, **(D)** GS15. Genes found in each gene module were used for the analysis. Each of the gene modules was enriched in the immune cluster. Top terms are shown.

**fig. S9. Correlation of metacells found in this study and metacells identified by Cole *et.al*** (***40***). **(A)** Metacells identified by cole *et al.* (n=125) were labelled according to cell-types (rows), metacells found in this study (n=187) are shown as columns. Pearson correlation coefficient is shown were red represents high correlation and blue represents low correlation.

**fig. S10. Transcription factors expression per metacell. (A)** Heatmap showing transcription factors expression across metacells. The dashed rectangle represents metacells that were enriched upon poly(I:C) treatment.

**Table S1. ImageStream raw** data used in **Fig. 1**.

**Table S2. DESeq2 results of RLRb high versus RLRb low immunostaining shown in** **Fig. 1F**.

**Table S3. GSEA results shown in** **Fig. 1G**.

**Table S4. DESeq2 results of mCherry positive versus mCherry negative sorted cells. Used in** **Fig. 3** **H-j.**

**Table S5. GSEA results shown in** **Fig. 3J**.

**Table S6. UMI fraction per metacell. Used for** **Fig. 5B**.

**Table S7. Differential expression analysis of poly(I:C) versus NaCl treated embryos using MAST on scRNAseq data.**

**Table S8**. Marker genes reported by Cole et al. (*40*)

**Table S9. Cell cluster markers identified in this study. Used in fig. S6. Table S10. MAST differential expression for cluster 1 and GSEA results. Table S11. WGCNA analysis for identifying gene modules (fig. S7) Table S12. ORA results for GS17 (brown) gene module.**

**Table S13. ORA results for GS16 (lightcyan) gene module. Table S14. ORA results for GS14 (red) gene module.**

**Table S15. ORA results for GS15 (purple) gene module.**

**Table S16. Marker genes identified in sub-clusters 0-3. Used in** **Fig. 6. Table S17. ORA results for sub-cluster 0.**

**Table S18. ORA results for sub-cluster 1. Table S19. ORA results for sub-cluster 2. Table S20. ORA results for sub-cluster 3.**

**Table S21. Orthofinder output used for orthogroups identification in** **Fig. 7**.

**Table S22. Shared single orthogroups and their gene expression used for correlation analysis in** **Fig. 7D** **Table S23. Shared genes with adjusted p value < 0.05 used for correlation analysis in** **Fig. 7E**.

Table S24. Genes that were shared between cluster in the non-injected control and the immune cluster in adult *Nematostella* identified by Cole et al. (*40*).

Table S25. **Dictionary for converting NVE gene models, NV2, and the gene models used in this study and their associated annotations.**

