## Supplementary Figures for "Functional characterization of specialized immune cells in a cnidarian reveals an ancestral antiviral program"

### Supplementary fig. 1

A

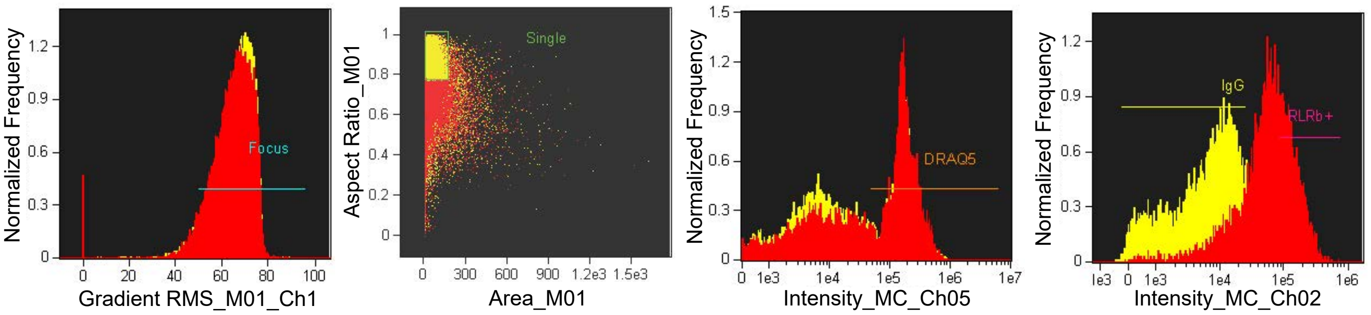


B

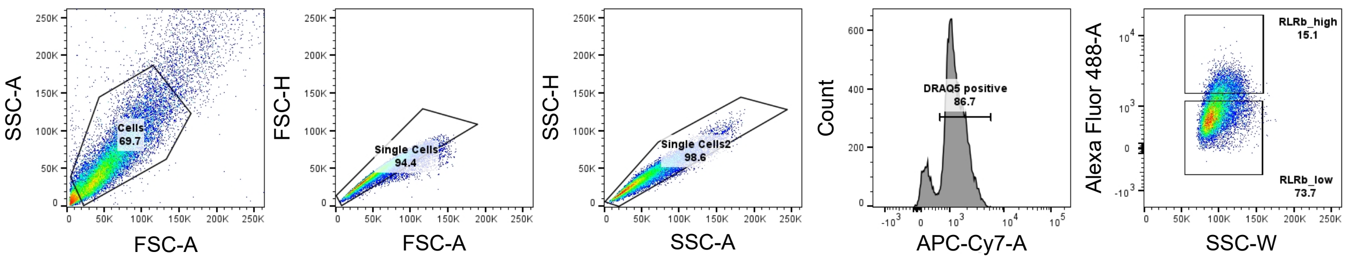

### Supplementary fig. 2

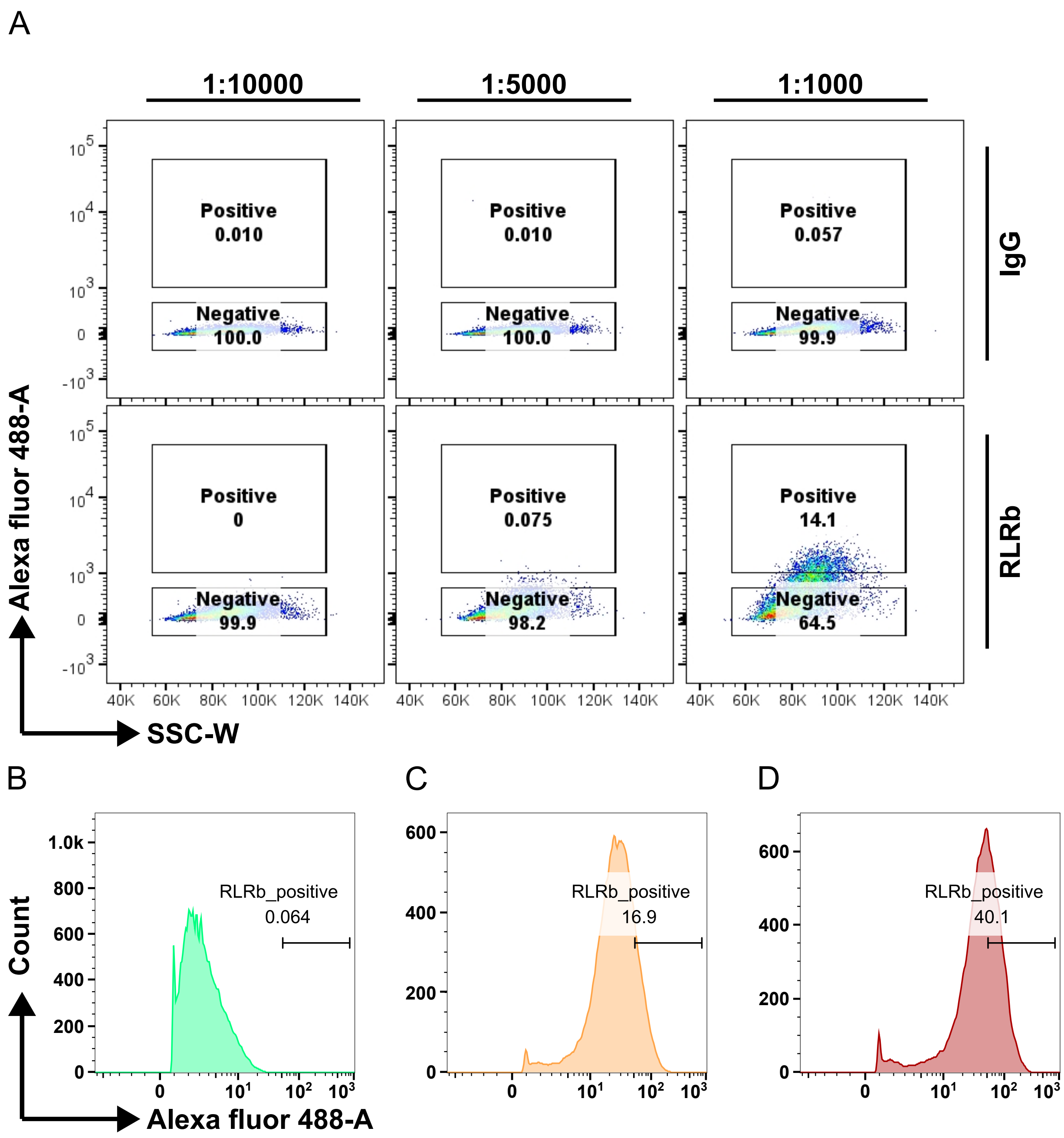

### Supplementary fig. 3

A

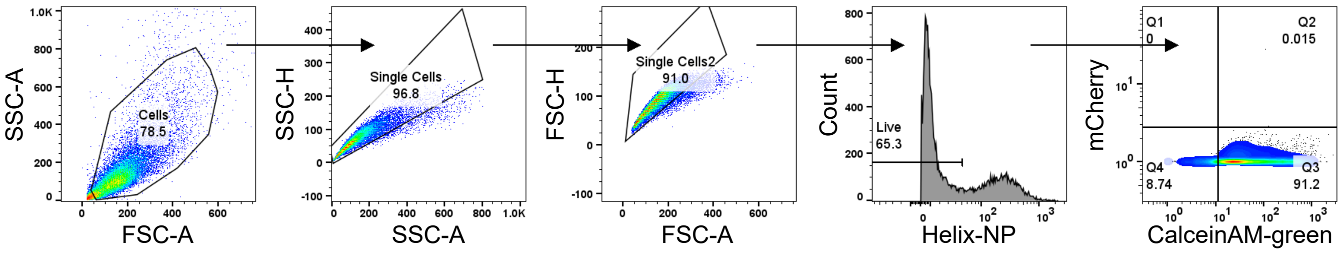

### Supplementary fig. 4

A

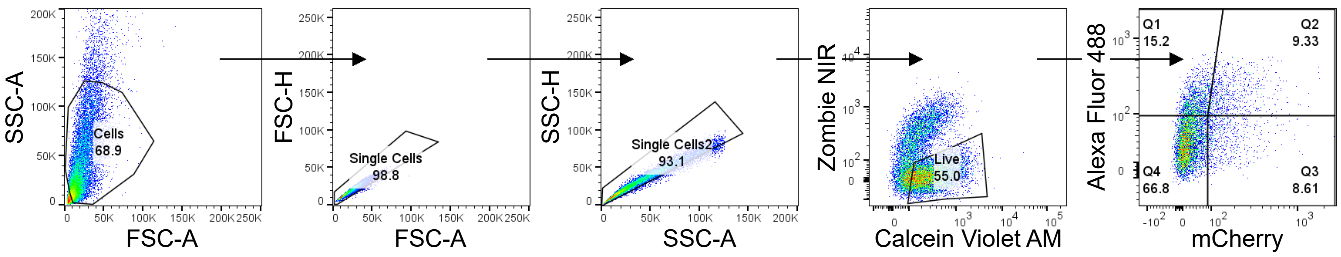

### Supplementary fig. 5

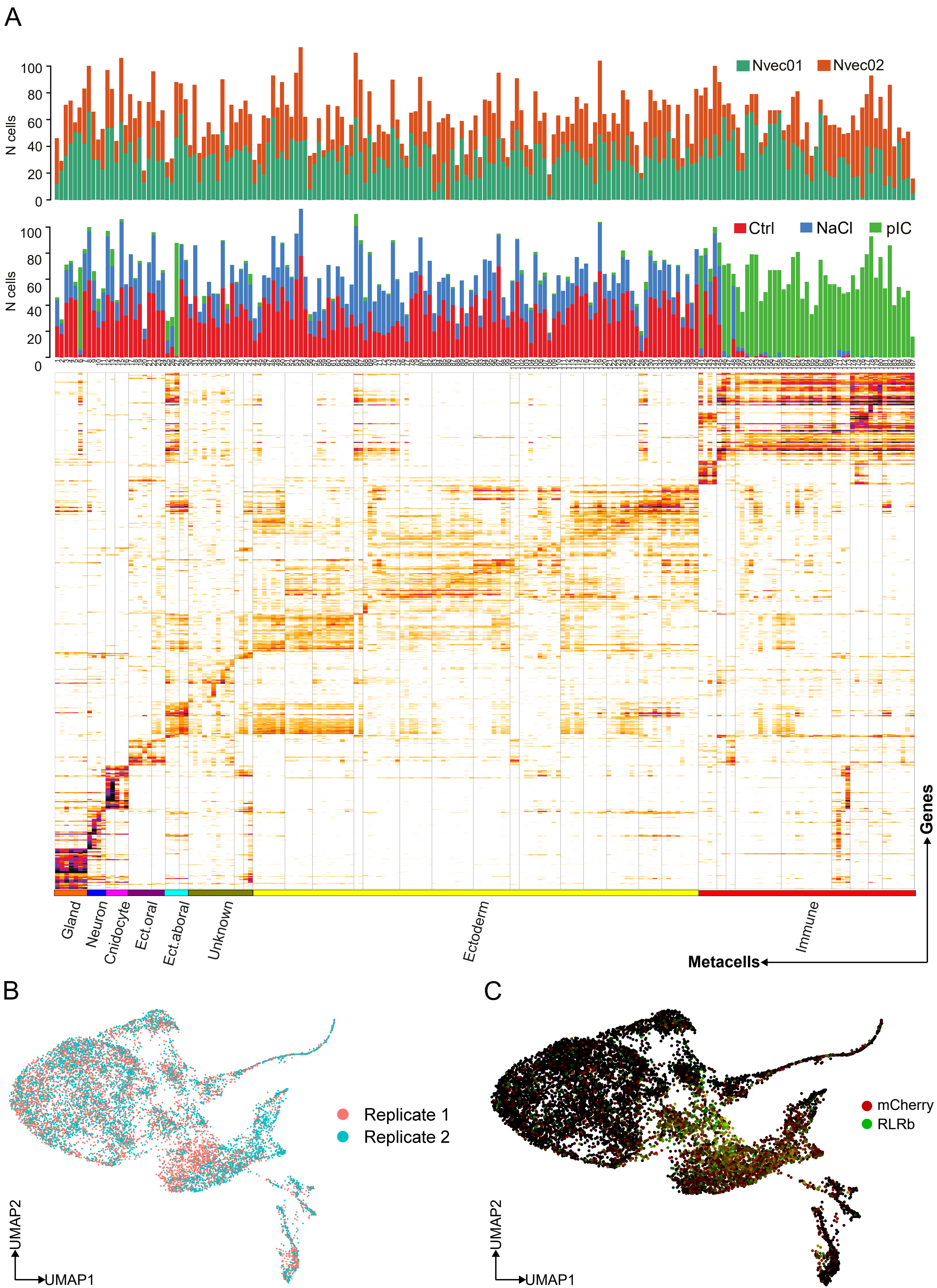

### Supplementary fig. 6

A

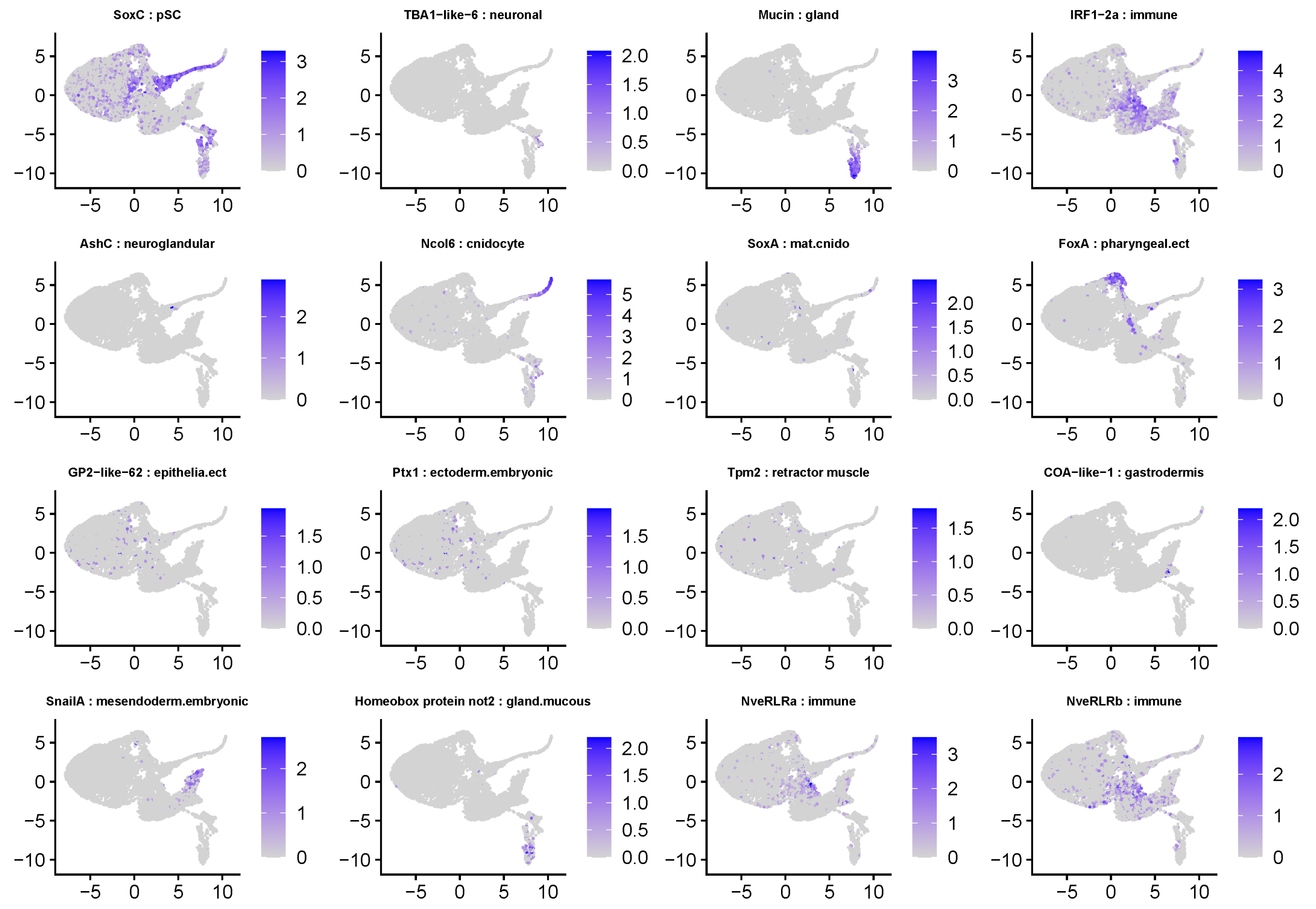

### Supplementary fig. 7

A

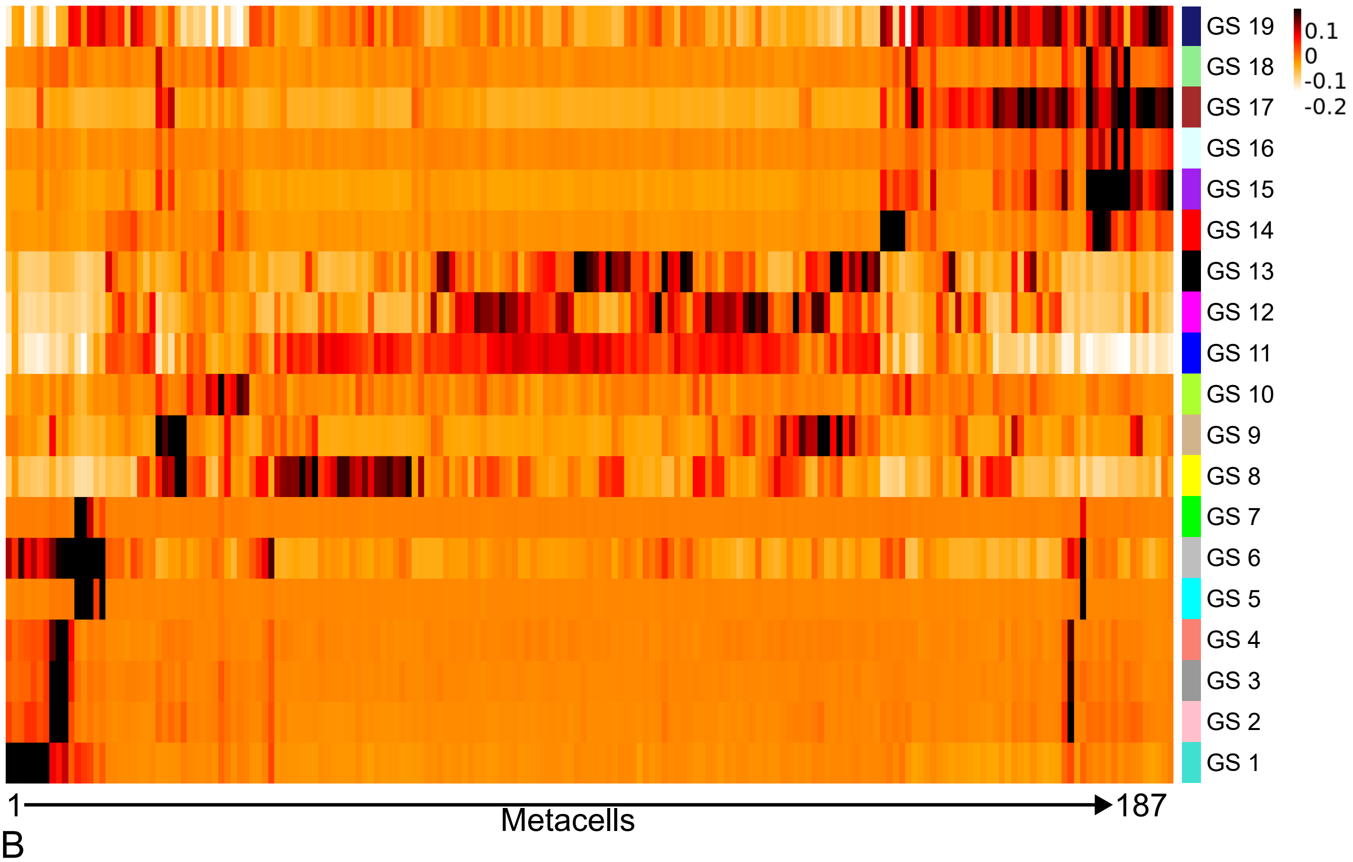


B

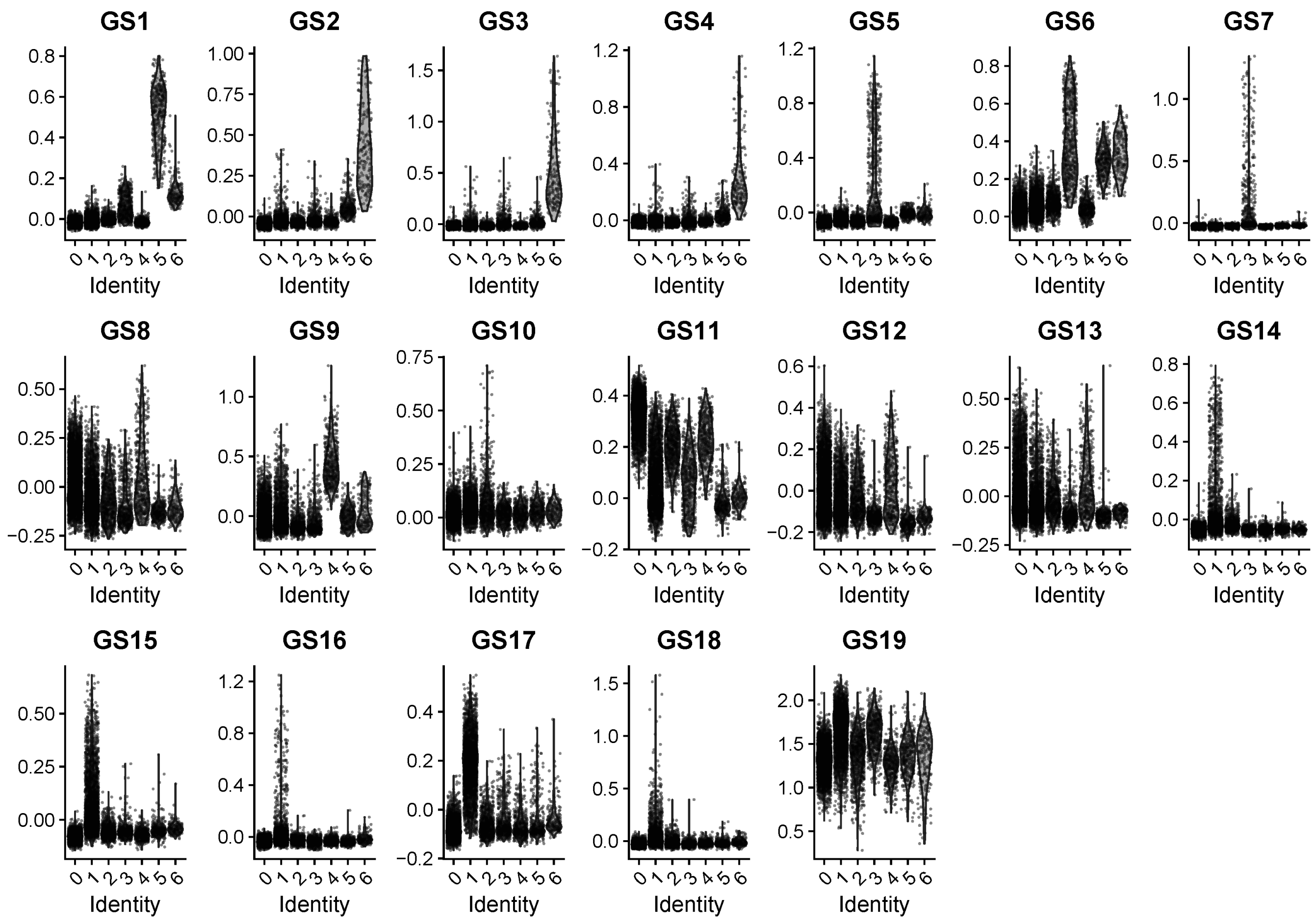

### Supplementary fig. 8

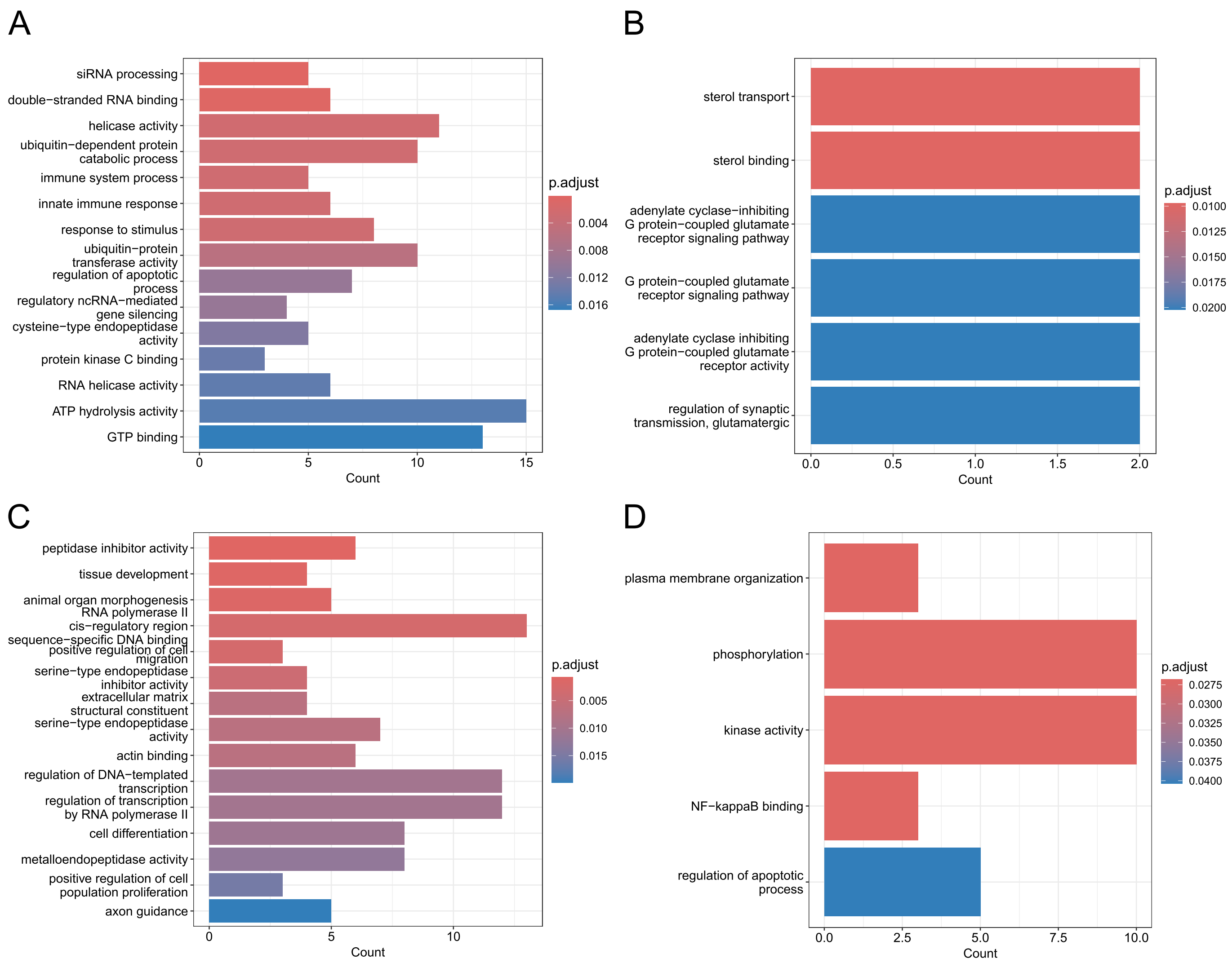

### Supplementary fig. 9

A

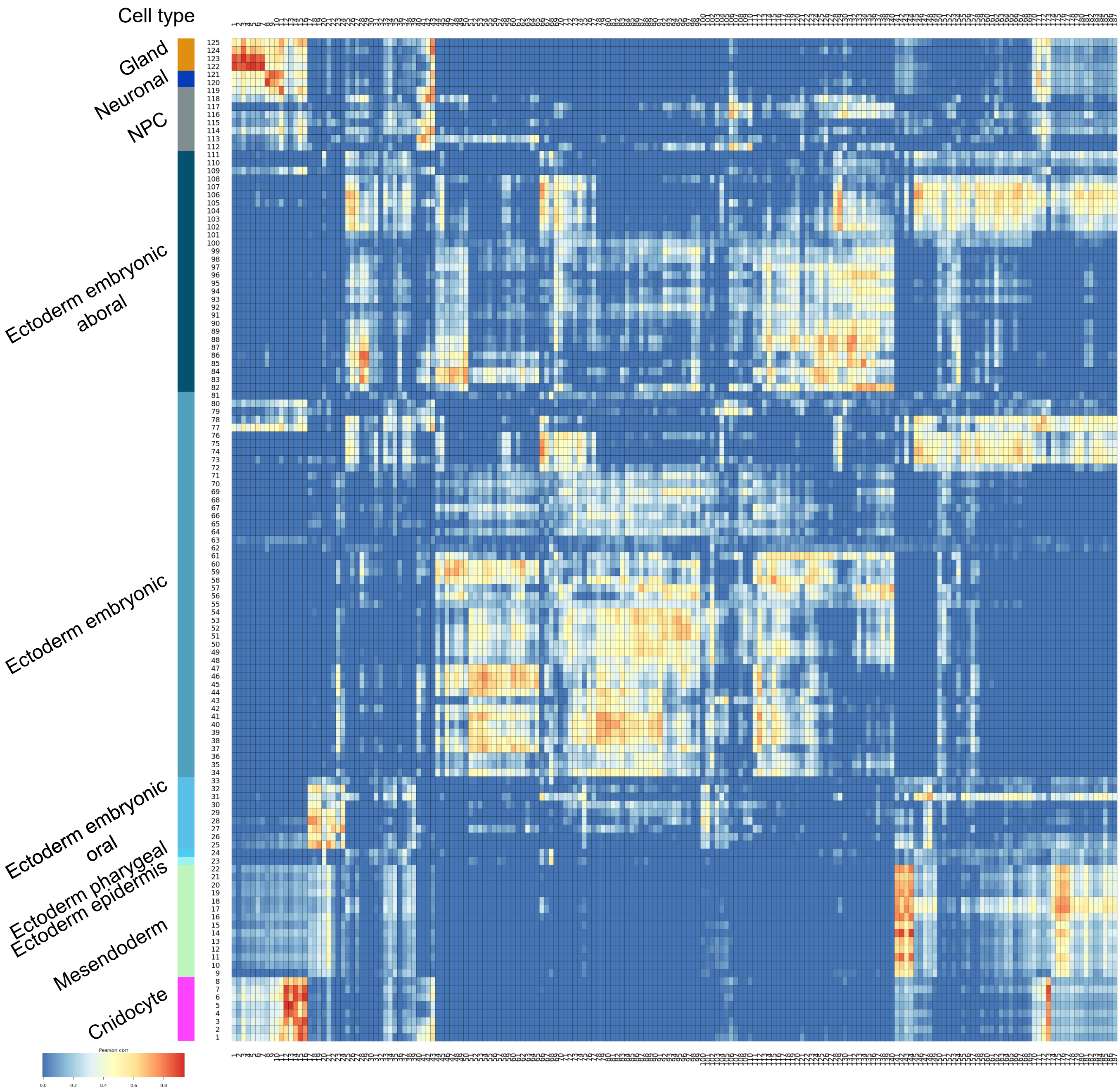

### Supplementary fig. 10

A

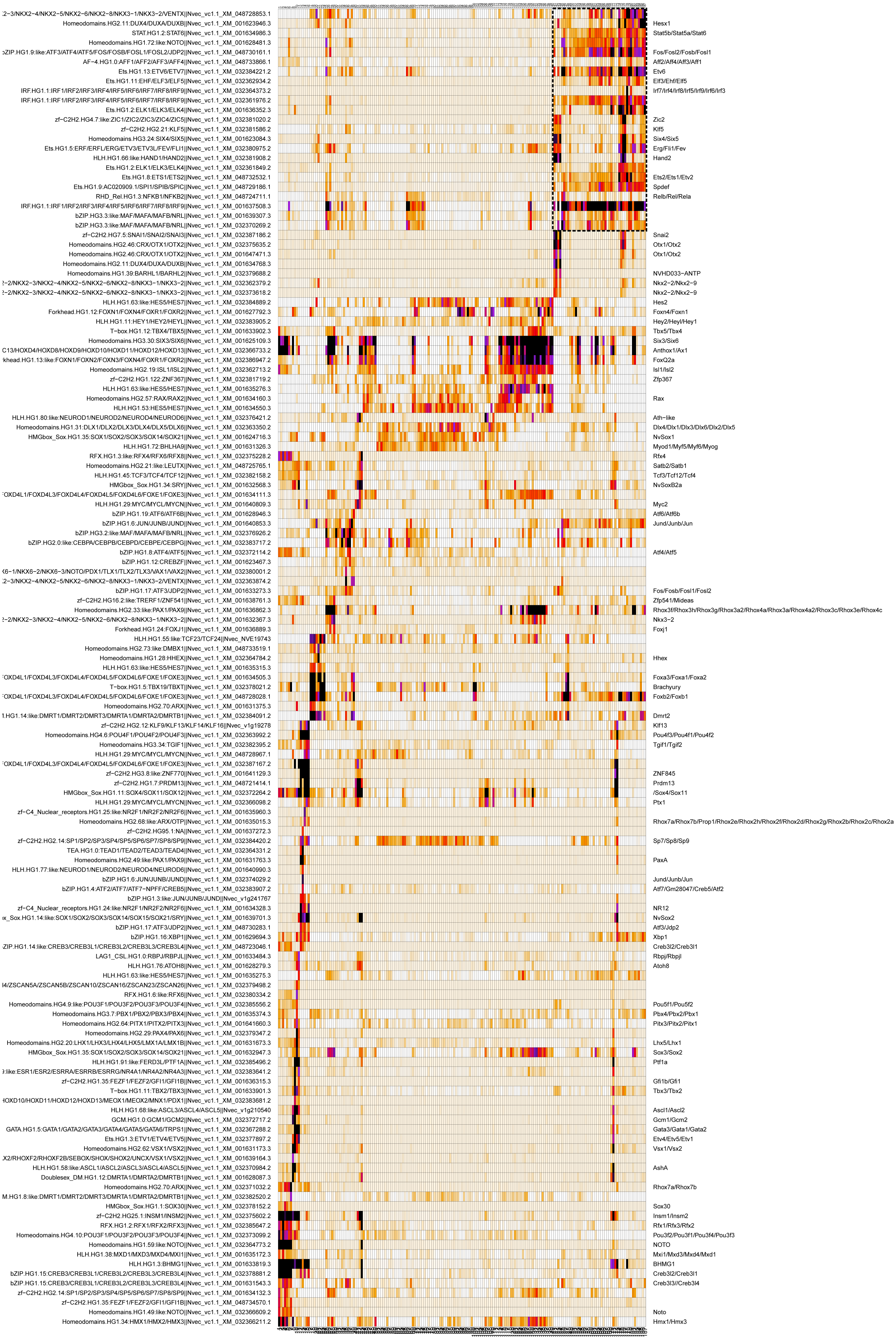
